# Self-reactive human CD8^+^ T cells restricted by HLA-E mediate immunoregulatory and antiviral functions

**DOI:** 10.64898/2026.10.12.758648

**Authors:** Lucy C. Walters, Xuan Guo, Carlos Casquero, Jonathan M. Urbach, Owen Leddy, Elif Ҫakan, Amanda O. Chan, Pallavi A. Balivada, Hannah Wisner, Bryan D. Bryson, Forest M. White, Mary Carrington, David R. Collins, Michael E. Birnbaum, Bruce D. Walker

## Abstract

HLA-E constitutively presents self VL9 peptides on healthy cells, signaling an intact HLA class I pathway and protecting cells from natural killer cell attack. Beyond this canonical role, HLA-E-restricted CD8⁺ T cells that recognize foreign pathogen peptides have emerged as mediators of immunity in multiple human infections, with orthologous responses conferring unprecedented protection in an HIV vaccine model. However, the poor stability of most pathogen-derived HLA-E ligands raises the possibility that additional modes of HLA-E-restricted recognition remain undiscovered. Here, we establish an alternative paradigm in which HLA-E-restricted CD8^+^ T cells recognize disease-associated increases in self-VL9/HLA-E rather than foreign antigen. These cells, which we term E-sensors, integrate activating and inhibitory signals to discriminate physiological from pathological levels of HLA-E expression. We demonstrate that E-sensors selectively eliminate highly activated, proliferating immune cells that express high levels of HLA-E, establishing a mechanistically resolved pathway of human CD8^+^ T cell immunoregulation. E-sensors also recognize pathogen-induced HLA-E upregulation and directly eliminate infected cells without prior pathogen-specific priming, demonstrating immediate immunosurveillance through recognition of increased self. Consistent with this antiviral function, we establish E-sensors as a previously unrecognized component of spontaneous HIV immune control in persons who naturally control infection. Together, our findings establish a pathway of human immunosurveillance and immunoregulation in which CD8⁺ T cells detect pathological increases in a constitutively expressed non-classical HLA class I molecule presenting a conserved self-peptide. This disease-agnostic mechanism of immune recognition has implications across infection, cancer, autoimmune disease, and other pathologies that upregulate HLA-E. Unlike classical HLA class I molecules, the near-monomorphic nature of HLA-E further positions this pathway as an untapped effector axis, warranting investigation as a potential target for immune interventions.

## INTRODUCTION

The role of HLA-E in protecting healthy cells from NK-cell attack is well established: by presenting VL9 peptides derived from classical HLA class I molecules, HLA-E signals that a cell has an intact HLA class I pathway and therefore should not be killed^1–5^. More recently, a new role for HLA-E in T cell biology has emerged. Whereas conventional CD8^+^ T cells express T cell receptors (TCR) that recognize foreign peptides presented by classical HLA-A, -B, and -C, unconventional CD8^+^ T cells whose TCRs instead recognize non-classical HLA-E are found in many human infections, including HIV^6,7^, *Mycobacterium tuberculosis*^8–10^, Epstein-Barr virus^11,12^, *Salmonella typhi*^13^, SARS-CoV-2^14^, hepatitis viruses^15–17^, and influenza^18^. Notably, orthologous MHC-E-restricted responses mediated remarkable protection in an HIV vaccine model^19–25^. Yet despite the apparent biological importance of HLA-E-restricted CD8^+^ T cells, their mechanistic basis remains poorly defined. Specifically, most pathogen-derived ligands proposed to bind HLA-E lack the sequence motifs required for HLA-E stabilization and therefore exhibit unprecedented instability^26–28^. This instability not only raises questions about the molecular basis of proposed HLA-E pathogen ligands but has also impeded therapeutic development^28^. It is therefore important to investigate whether alternative mechanisms could contribute to HLA-E-restricted CD8^+^ T cell immunity.

Unstable pathogen peptide presentation by HLA-E is sharply contrasted by the stable presentation of VL9^26,29^. Structurally, the HLA-E binding groove is highly specialized and forms extensive contacts with VL9 peptides, which limits the sequence variability of peptides that can stably bind HLA-E^26,29,30^ and results in a VL9-dominated peptide repertoire^5,28^. Healthy cells constitutively express HLA-E/VL9, which inhibits NK cell attack. However, in disease, HLA-E/VL9 is often either down- or up-regulated. Pathogens and tumors frequently disrupt the HLA class I pathway to evade conventional CD8^+^ T cells; however, this limits the supply of HLA-A/B/C-derived VL9 peptides and results in HLA-E downregulation^31,32^. Many pathogens and tumors counter this by upregulating HLA-E to preserve NK cell inhibition and avoid elimination^33–35^. Human cytomegalovirus (HCMV) provides a well-characterized example of this: by encoding its own VL9 peptide that closely mimics human VL9 sequences, HCMV upregulates HLA-E/VL9 and evades NK attack despite disrupting classical HLA class I antigen presentation^35,36^.

HLA-E presentation of HCMV-derived VL9 also results in the induction of HLA-E-restricted VL9-specific CD8⁺ T cells^37,38^. In prior studies, higher-affinity interactions with HCMV-derived VL9 than with self-VL9, combined with co-expression of inhibitory KIR receptors, prevented self-reactivity and enabled HCMV-specific responses^37^. In an unrelated study, KIR^+^ CD8^+^ T cells were shown to mediate immunoregulatory functions and prevent autoimmune disease, although the mechanistic basis was not elucidated^39^.

Given these findings, we asked whether HLA-E-restricted CD8⁺ T cells can sense disease-associated increases in self-VL9/HLA-E rather than foreign antigen. Under healthy conditions, lower HLA-E expression and dominant inhibitory KIR2DL–HLA-C signaling should restrain activation of self-VL9-reactive HLA-E-restricted CD8⁺ T cells^37^. In contrast, many viruses and cancers upregulate HLA-E to maintain inhibition of CD94/NKG2A^+^ NK- and T-cells^33–35^, while inflammation, immune activation, and cellular stress can also enhance HLA-E expression^40–43^–hypothetically increasing signaling through activating self-VL9/HLA-E-specific TCRs and shifting the balance from self-tolerance towards immunosurveillance. In parallel, pathogen-mediated HLA-C downregulation^44^ and stress-associated remodeling of the HLA-C peptidome can weaken inhibitory KIR engagement^45^ which may further potentiate HLA-E/VL9-specific CD8^+^ T cells. We therefore hypothesized that self-VL9-reactive HLA-E-restricted CD8⁺ T cells may be uniquely positioned to exploit disease-associated HLA-E upregulation as an activating cue, transforming this conserved mechanism of immune evasion and inhibition into a substrate for immune detection. Under this model, pathological HLA-E upregulation is not merely a brake on immunity but a signal that licenses HLA-E/VL9-specific CD8⁺ T cells, which we term E-sensor T cells (E-sensors), enabling direct, disease-non-specific immunosurveillance.

## RESULTS

### SIV/HIV MHC-E epitopes associated with vaccine protection do not measurably stabilize MHC-E

MHC-E-restricted CD8^+^ T cell responses mediated remarkable protection in a preclinical HIV vaccine model in rhesus macaques^25^. Notably, these responses were reported to recognize SIV-derived epitopes that lack canonical MHC-E-binding sequence motifs, raising questions about the molecular basis of their presentation and recognition^19^. We therefore performed thermostability assays to directly assess the capacity of proposed vaccine-associated SIV epitopes and their aligned HIV counterparts to bind rhesus Mamu-E and human HLA-E. Eight out of nine vaccine-identified SIV epitopes and their corresponding HIV counterparts showed no measurable binding to Mamu-E or HLA-E, as indicated by no increase in thermostability above peptide-free baselines **(Fig. 1a)**. For these eight non-binders, we measured Mamu-E and HLA-E stability in the presence of increasing concentrations of supraphysiologic SIV/HIV peptide but detected no peptide-induced stabilization even at the highest peptide concentration. Only one vaccine-identified SIV epitope, RL9SIV (sequence: RMYNPTNIL), and its HIV counterpart peptide, RL9HIV (sequence: RMYSPTSIL), demonstrated measurable MHC-E stabilization **(Fig 1a)**. However, despite using supraphysiologic peptide concentrations and obtaining measurements in the presence of excess peptide with no prior wash steps, RL9HIV/HLA-E and RL9SIV/Mamu-E exhibited low thermal melting temperatures (T_m_s) around or below physiological temperature (37°C), raising questions about whether these peptides can be stably presented *in vivo*. In sharp contrast, positive control VL9 peptides robustly stabilized both human and rhesus MHC-E orthologs, yielding T_m_s that far exceeded physiologic temperature across all tested peptide concentrations.

**Figure 1.**
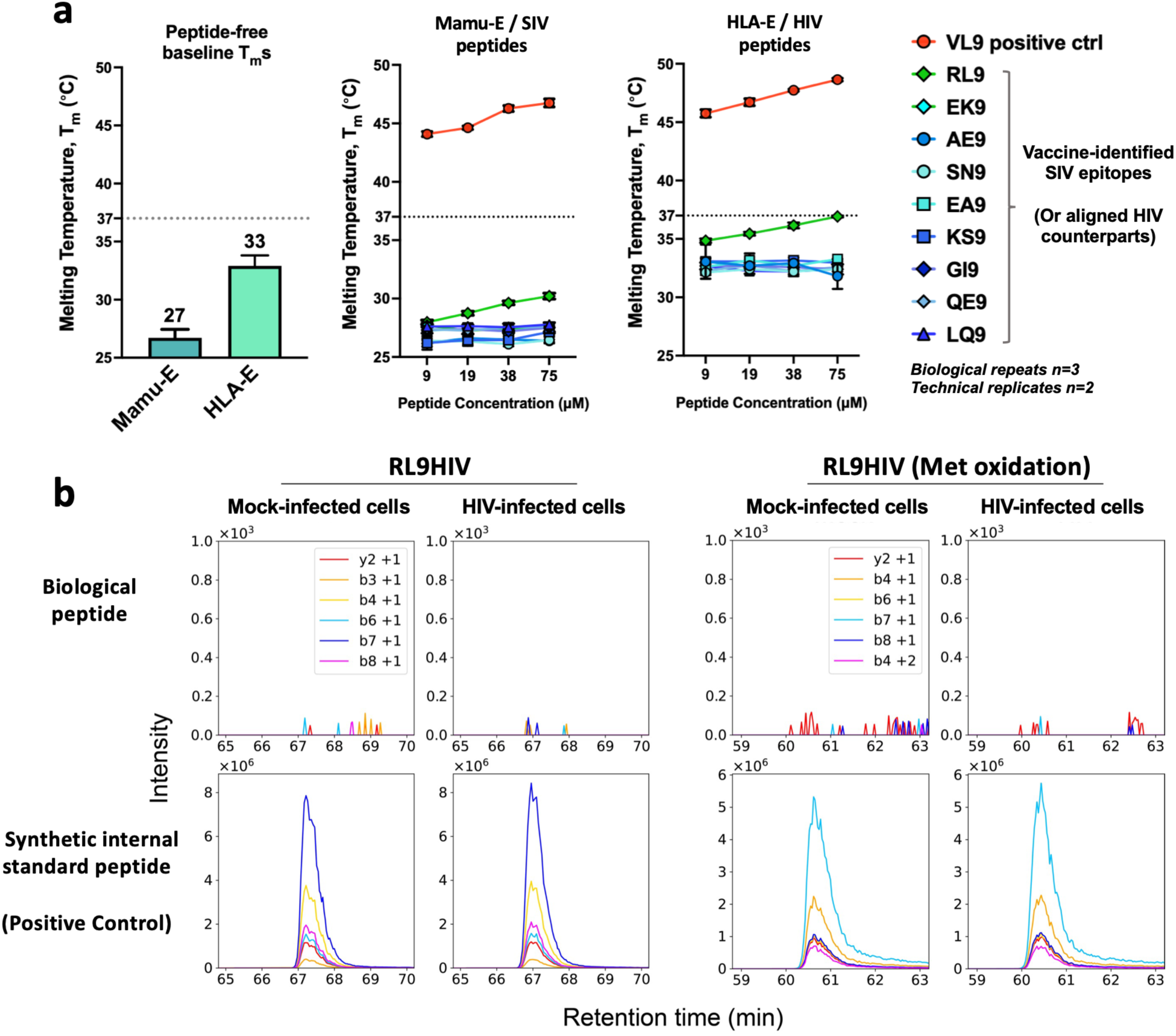
Vaccine-identified MHC-E epitopes derived from SIV / HIV do not stably bind human or rhesus MHC-E. **A.** Left panel shows thermostability of human HLA-E and rhesus Mamu-E orthologs pre-refolded with human or rhesus β2-microglobulin in the absence of any added peptide. Mean thermal melting temperatures (T_m_s) of ‘empty’ HLA-E and Mamu-E complexes are indicated above bars, and error bars indicate standard deviation from five independent biological repeats. The dashed line at 37°C indicates normal physiologic temperature for rhesus macaques and humans. The middle panel shows thermostability of Mamu-E following incubation with a positive control VL9 peptide (VMAPRTLVL) or vaccine-identified MHC-E epitopes derived from SIV (peptide sequences are listed in the methods). Peptide-induced stability is measured at increasing concentrations of test peptide. Error bars indicate standard deviation from three independent biological repeats. The right panel is identical to the middle panel except that human HLA-E is shown, and the peptides are HIV-derived sequence-aligned counterparts of vaccine-identified SIV epitopes. **B.** Representative extracted ion chromatograms from SureQuant mass spectrometry analysis of HLA-E-bound peptides isolated from mock- or HIV-infected cells (HLA-E-transfected K562 cell lines, pseudotyped NL4-3 HIV virus). Heavy stable isotope-labeled (SIL) RL9HIV peptides added to each sample were readily detected (bottom panels), validating assay performance and expected retention times. In contrast, no endogenous (light) RL9HIV peptide signal was detected in HLA-E immunopeptidomes from either mock- or HIV-infected cells (top panels) in either its unmodified or oxidized form.

Given its low thermostability, we investigated whether the RL9HIV peptide can be endogenously presented by HLA-E on HIV-infected cells. However, despite overexpressing HLA-E and using targeted mass spectrometry searches sensitive to as little as 1 molecule per cell of each target peptide-MHC complex^46,47^, RL9HIV was not detected **(Fig. 1b)**. These data are consistent with prior studies in which unstable RL9HIV presentation impeded HLA-E-based therapeutic development^28^. In contrast to RL9HIV, VL9 peptides were readily detectable by mass spectrometry in both HIV-infected and mock-infected cells **(Extended Data Fig. 1)**.

### Persistent and highly expanded E-sensor populations are NK-like and exceptionally cytotoxic

Given that many reported pathogen-derived HLA-E epitopes bind HLA-E weakly or not at all^27,28,48^, we reasoned that additional modes of HLA-E-restricted T cell recognition may exist. We therefore asked whether HLA-E-restricted CD8^+^ T cells can operate by sensing disease-associated increases in HLA-E presenting its canonical self-peptide, VL9. To identify donors with HLA-E-restricted VL9-specific CD8⁺ T cell responses, we assembled a cohort of 28 healthy individuals selected for characteristics previously associated with an increased likelihood of harboring these responses, including HCMV seropositivity and particular HLA class I allelic backgrounds **(Extended Data Table 1)**^37,38^. Accordingly, this cohort was not intended to estimate the prevalence of these responses in the general CMV-seropositive population, but rather to maximize the likelihood of identifying individuals with these responses for downstream phenotypic and functional analyses.

We performed direct *ex vivo* HLA-E/VL9 tetramer staining and incorporated anti-CD94 blockade to prevent tetramer binding to NKG2 receptors **(Extended Data Fig. 2)**, thereby isolating TCR-dependent tetramer staining on HLA-E-restricted VL9-specific CD8^+^ T cells^37^. Consistent with prior reports^37,38^, HLA-E/VL9 tetramer^+^ E-sensors were readily detected, with 10 of 28 HCMV-seropositive donors exhibiting highly expanded and longitudinally persistent populations ranging from 1% to 15.5% of the CD8^+^ T cell repertoire **(Fig. 2a-c)**. Additional donors exhibited smaller but well-resolved E-sensor populations, such as 0.2% of CD8^+^ T cells in Donor 12, demonstrating that the size of the response can be highly variable across individuals **(Fig. 2a).** No cross-recognition by HLA-E tetramers presenting previously reported *Mycobacterium tuberculosis*^49^ or SARS-CoV-2^14^ HLA-E epitopes was observed, supporting VL9 peptide specificity **(Fig. 2d)**. Equally, no resolvable HLA-E/VL9-specific CD8^+^ T cell responses were detected in CMV-seronegative donors enriched for the same characteristics, consistent with previous reports identifying these responses exclusively in CMV-seropositive individuals.^37,38^ **(Fig. 2b) (Extended Data Table 2)**.

**Figure 2.**
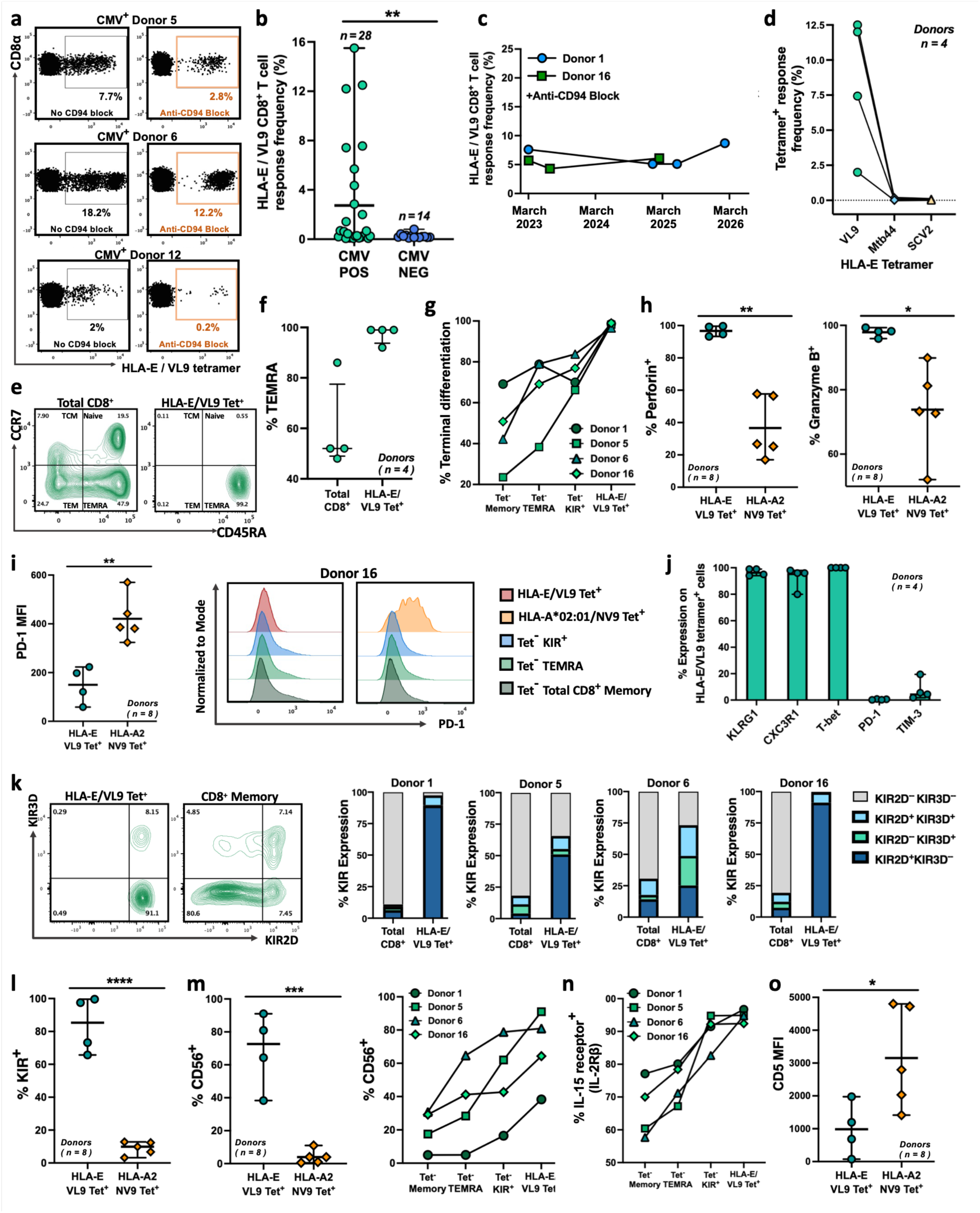
Highly expanded and persistent E-sensor populations exhibit a cytotoxic effector-ready phenotype with NK-like features. **(a)** Representative HLA-E*01:03/VL9 (VMAPRTLIL) tetramer staining with or without anti-CD94 blockade to isolate TCR-dependent tetramer binding. Plots are gated on live singlet CD3⁺CD8⁺ T cells. Three CMV⁺ donors with distinct, MFI-resolvable tetramer⁺ HLA-E-restricted VL9-specific CD8^+^ T cell populations are shown. **(b)** Frequency of HLA-E-restricted VL9-specific tetramer⁺ CD8⁺ T cells among total CD8⁺ T cells in 28 CMV⁺ and 14 CMV⁻ donors enriched for HLA class I backgrounds previously associated with HLA-E/VL9 responses. Horizontal lines indicate means; error bars indicate the range. Statistical significance was assessed using an unpaired two-tailed Welch’s t-test (P = 0.0057). **(c)** Longitudinal frequencies of HLA-E/VL9 tetramer⁺ CD8⁺ T cells in CMV⁺ donors. Lines connect donor-matched measurements across time points. **(d)** Frequency of HLA-E tetramer⁺ CD8⁺ T cells in four CMV⁺ donors stained with tetramers loaded with canonical VL9 (VMAPRTLIL), the *Mycobacterium tuberculosis*-derived peptide Mtb44 (RLPAKAPLL), or a previously reported SARS-CoV-2-derived HLA-E-binding peptide (VMPLSAPTL). Symbols represent individual donors; lines connect donor-matched measurements. **(e)** Representative CCR7 versus CD45RA staining of total and HLA-E/VL9 tetramer⁺ CD8⁺ T cells from Donor 16. **(f)** Frequency of TEMRA cells among total CD8⁺ T cells and HLA-E/VL9 tetramer⁺ CD8⁺ T cells from CMV⁺ Donors 1, 5, 6, and 16, who harbor highly expanded HLA-E/VL9 tetramer^+^ CD8^+^ T cell populations. **(g)** Frequency of terminally differentiated (CD27⁻CD28⁻) cells among HLA-E/VL9 tetramer⁺ CD8⁺ T cells and comparator tetramer-negative CD8⁺ T-cell subsets. **(h)** Frequency of perforin⁺ (left) and granzyme B⁺ (right) cells among HLA-E/VL9 tetramer⁺ and HLA-A*02:01/NV9 (an immunodominant HCMV-derived peptide) tetramer⁺ CD8⁺ T cells. **(i)** Left, comparison of PD-1 MFI between HLA-E/VL9 and HLA-A*02:01/NV9 tetramer⁺ CD8⁺ T cells. Right, representative PD-1 histograms from Donor 16. **(j)** Frequency of PD-1⁺, TIM-3⁺, KLRG1⁺, CX3CR1⁺, and T-bet⁺ cells among HLA-E-restricted VL9-specific tetramer⁺ CD8⁺ T cells. **(k)** Left, representative KIR2D versus KIR3D staining of HLA-E/VL9 tetramer⁺ and tetramer⁻ memory CD8⁺ T cells from Donor 16. Right, KIR2D and KIR3D expression patterns among HLA-E-restricted VL9-specific tetramer⁺ versus total CD8⁺ T cells. **(l)** Frequency of KIR-expressing cells among HLA-E/VL9 tetramer⁺ versus HLA-A*02:01/NV9 tetramer⁺ CD8⁺ T cells. **(m)** CD56 expression on HLA-E/VL9 tetramer⁺ versus HLA-A*02:01/NV9 tetramer⁺ CD8⁺ T cells (left) and donor-matched CD8⁺ T-cell subsets (right). **(n)** IL-15 receptor β chain (IL-2Rβ, CD122) expression on HLA-E-restricted VL9-specific tetramer⁺ and donor-matched tetramer⁻ CD8⁺ T-cell subsets. **(o)** CD5 (a molecule reported to dampen IL-15 signaling) MFI on HLA-E/VL9 tetramer⁺ versus HLA-A*02:01/NV9 tetramer⁺ CD8⁺ T cells. *Unless otherwise indicated, HLA-E/VL9 tetramer⁺ populations were identified under CD94 blockade. Phenotypic HLA-E/VL9 analyses were performed on four CMV⁺ healthy donors (D1, D5, D6, and D16), and HLA-A*02:01/NV9 analyses on four CMV⁺ healthy donors plus Donor 16, where the presence of tetramer^+^ responses was pre-validated. Symbols represent individual donors. Horizontal bars indicate means; error bars indicate the range. Statistical significance is indicated as follows: ns, P > 0.05; *P ≤ 0.05; **P ≤ 0.01; ***P ≤ 0.001; ****P ≤ 0.0001*.

TCR sequencing of E-sensors revealed significantly less diversity than bulk-sorted CD8^+^ T cells, but comparable levels to classically restricted CMV-specific CD8^+^ T cell responses **(Extended Data Fig. 3a)**. Conservative clonotypic filtering revealed that E-sensors exhibit a highly oligoclonal repertoire with evidence of partially convergent selection **(Table 1)**. For example, TRAJ7 usage was significantly enriched, and TRAV21-TRAJ7 pairing with length-matched CDR3α sequences was identified in independent donors. Consistent with prior reports of HLA-E/VL9-specific TCR architecture^37,50^, 11 of 17 clonotypes contained arginine residues within the CDR3β sequence. Together, these findings reveal structural features consistent with partially convergent selection within an otherwise donor-skewed oligoclonal HLA-E/VL9-specific TCR repertoire.

**Table 1.**
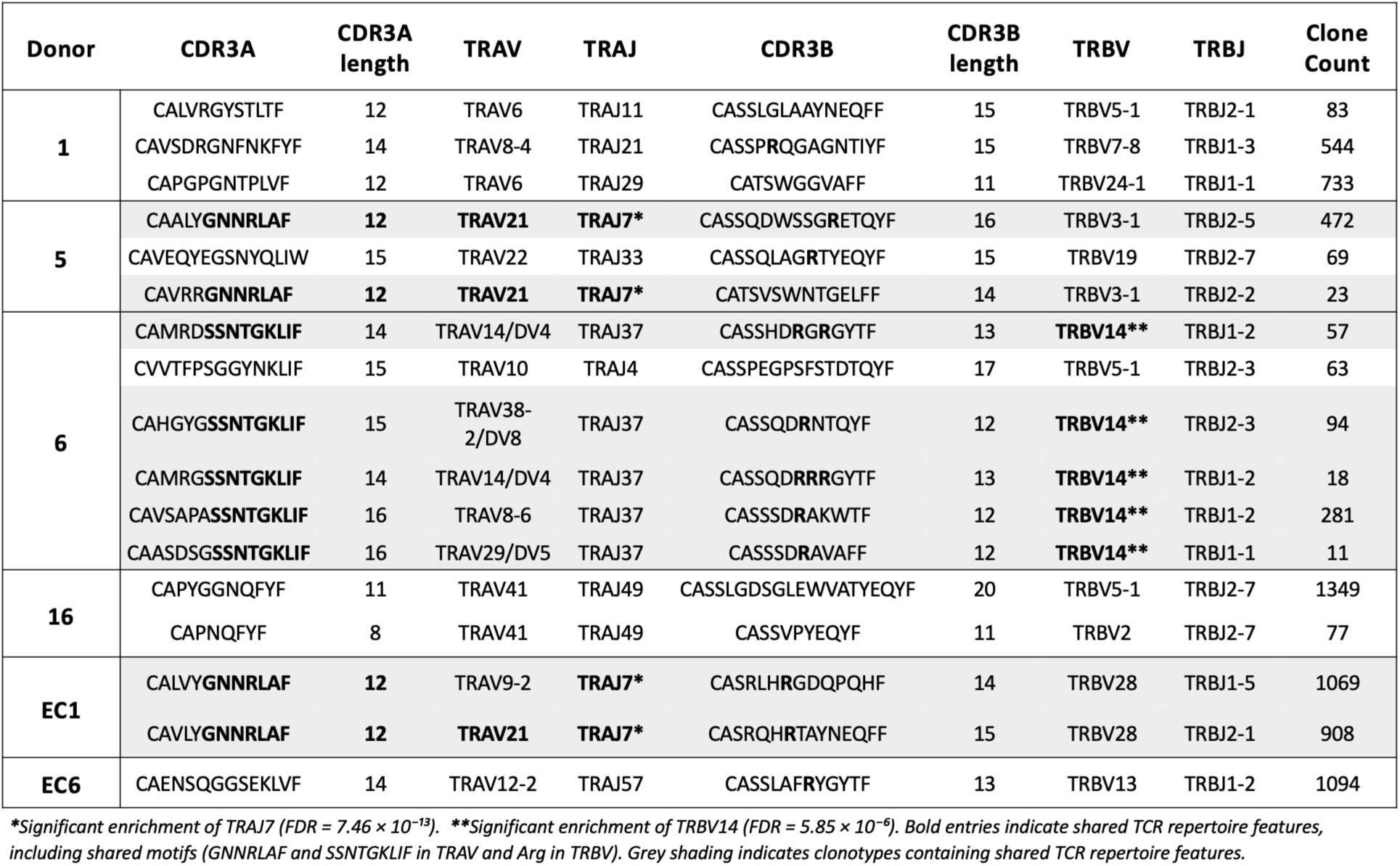
Enriched HLA-E-restricted VL9-specific TCR clonotypes. Paired TCRα and TCRβ sequences obtained by single-cell V(D)J sequencing of HLA-E*01:03/VL9 tetramer⁺ CD8⁺ T cells from CMV^+^ healthy donors (D1, D5, D6, and D16) and HIV elite controllers (EC1 and EC6). Cells were sorted as live single CD3⁺CD8⁺ tetramer⁺ lymphocytes following CD94 blockade. Clonotypes met conservative enrichment criteria of ≥10 cells and ≥2-fold enrichment relative to donor-matched bulk CD8⁺ T-cell repertoires.

Phenotypic profiling of E-sensors demonstrated advanced terminal differentiation, with near-complete loss of the costimulatory markers CD27 and CD28 (mean, 98% double-negative) and a predominant TEMRA phenotype (CD45RA⁺CCR7⁻; mean, 97%) **(Fig. 2e-g)**. E-sensors also expressed high CD57 relative to comparator subsets (including tetramer-negative KIR^+^, TEMRA, and memory CD8^+^ T cells) – a terminal effector marker previously linked to reduced proliferative capacity **(Extended Data Fig. 3b,c)**^51^. Despite this terminal phenotype, E-sensors retained a highly cytotoxic program, with near-universal *ex vivo* expression of granzyme B and perforin (mean, 98% and 97%, respectively), exceeding all donor-matched CD8^+^ T cell comparator populations **(Extended Data Fig. 2d)** and significantly surpassing comparator HLA-A2-restricted CMV-specific CD8^+^ T cells **(Fig. 2h)**. E-sensors exhibited minimal expression of the negative immunoregulatory molecules PD-1 and TIM-3, alongside uniformly high T-bet, KLRG1, and CX3CR1 expression, indicative of a non-exhausted effector-like program **(Fig. 2i,j) (Extended Data Fig. 3e)**. Exclusive expression of αβ TCRs further distinguished E-sensors from the broader KIR⁺ CD8⁺ pool, which included both αβ and γδ subsets **(Extended Data Fig. 3f)**.

E-sensors also displayed a distinctive NK receptor-rich phenotype. KIR2D expression predominated, with half the donors exhibiting near-uniform E-sensor KIR expression (>95%) **(Fig. 2k)**. Within the total KIR^+^ CD8^+^ pool, E-sensors comprised a substantial portion, with 19-35% of KIR^+^ CD8^+^ T cells staining positive for the HLA-E/VL9 tetramer **(Extended Data Fig. 3g)**. E-sensors also expressed significantly higher CD56, an NK cell-associated hallmark of potent cytotoxicity, compared to HLA-A2/CMV-specific comparators, and 2–7-fold higher CD56 levels than donor-matched bulk CD8^+^ memory **(Fig. 2m)**. By contrast, CD16 expression, an NK-associated marker linked to TCR-independent function in CD8^+^ T cells^52^, did not differ significantly from HLA-A2/CMV-specific cells **(Extended Data Fig. 3h)**. KIR⁺ CD8⁺ T cells lacking HLA-E/VL9 specificity exhibited the highest CD16 expression, whereas E-sensors showed 2–9-fold lower levels, indicating E-sensors may be more TCR-dependent than other KIR^+^ CD8^+^ T cells **(Extended Data Fig. 3i)**. E-sensors also exhibited near-uniform IL-15 receptor β chain (IL-2Rβ) expression **(Fig. 2n)** and significantly reduced MFI (up to 53-fold) of CD5–which can dampen IL-15-signaling^53^–relative to HLA-A2/CMV-specific cells and CD8^+^ comparator populations, consistent with heightened IL-15 sensitivity **(Fig. 2o)**. Collectively, these data define E-sensors as a potently cytotoxic HLA-E-restricted, innate-like terminal effector memory lineage that is phenotypically distinct from classical CMV-specific responses, the broader KIR⁺ CD8⁺ pool, and canonically exhausted states.

### ‘Increased self’ licenses E-sensors while inflammatory cues further amplify the response

In order for HLA-E/VL9-specific CD8⁺ T cells to mediate immunosurveillance by sensing disease-associated HLA-E upregulation, beyond recognizing the HCMV UL40-derived VL9 mimetic, these cells would need to recognize HLA-E presenting endogenous self-VL9 peptide. We demonstrate strong self-VL9 recognition by E-sensors across all tested donors, with 3-11% of the CD8^+^ T cell repertoire staining positively for donor-matched self-VL9/HLA-E tetramer **(Fig. 3a)**. We next tested whether this self-reactivity enables E-sensors to break tolerance and respond to disease-associated increases in HLA-E. Increased HLA-E expression was modeled by stimulating PBMCs with high concentrations of tetrameric HLA-E complexes presenting donor-matched self-VL9. In our model system, HLA-E upregulation indeed broke tolerance and induced modest yet selective activation and degranulation of E-sensors, while bulk (E-sensor-negative) CD8⁺ T cells remained unresponsive (**Fig. 3b**). The level of E-sensor activation scaled with the MFI of the HLA-E/VL9 tetramer used to model HLA-E upregulation: HLA-E tetramers loaded with VL9 variants that stained with higher MFI, consistent with stronger TCR–HLA-E/VL9 interactions, also elicited stronger effector activity when used to model HLA-E upregulation **(Fig. 3c,d)**.

**Figure 3.**
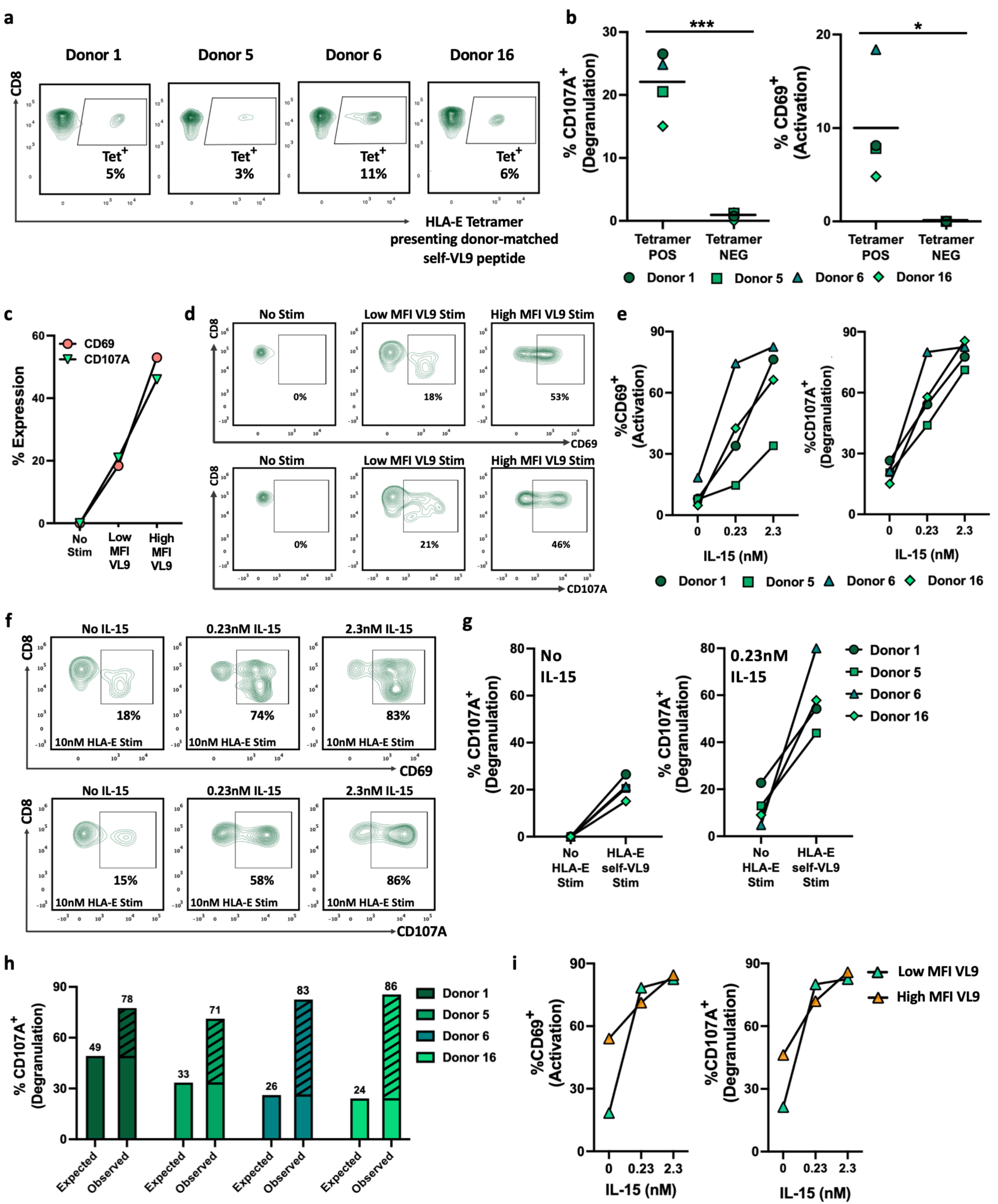
E-sensors detect increased HLA-E, and IL-15 further amplifies effector activity in the absence of foreign antigen. **(a)** Representative staining by HLA-E tetramers presenting donor-matched self-VL9 peptides in four CMV⁺ donors with expanded E-sensor populations. **(b)** CD107a (degranulation, left) and CD69 (activation, right) expression in HLA-E/VL9 tetramer⁺ and tetramer⁻ CD3^+^CD8⁺ T cells following stimulation with donor-matched self-VL9/HLA-E tetramers. Each dot represents an independent donor; horizontal lines indicate group means. **(c)** Plot showing data from d**. (d)** CD69 and CD107a expression on E-sensors from Donor 6 following stimulation with HLA-E tetramers presenting different VL9 peptides that yield either low- or high-MFI populations by tetramer staining. Plots are gated on live singlet CD3^+^CD8^+^ HLA-E/VL9 tetramer^+^ T cells; gates indicate CD107A^+^ or CD69^+^ cells. **(e)** Plots summarizing data in f across four donors. The percentage of E-sensors expressing CD69^+^ or CD107A^+^ is plotted on the Y-axis. **(f)** Representative Donor 6 CD69 and CD107A expression following stimulation with donor-matched self-VL9/HLA-E tetramers with and without increasing concentrations of IL-15 superagonist. Plots are gated as in d. **(g)** The percentage of E-sensors expressing CD107A with or without donor-matched self-VL9/HLA-E tetramer stimulation in the absence (left) or presence (right) of preincubation with an IL-15-based superagonist. **(h)** The percentage of E-sensors expressing CD107A following stimulation with donor-matched self-VL9/HLA-E tetramers, IL-15 superagonist, or both. Expected additive responses were calculated from individual stimuli and compared with observed responses following combined stimulation. Dashed regions indicate synergistic degranulation exceeding the predicted additive response. **(i)** CD69 (left) and CD107a (right) expression in Donor 6 following stimulation with HLA-E tetramers loaded with VL9 variants that stained with low or high MFI across increasing IL-15 superagonist concentrations. *Unless otherwise indicated, analyses were performed on HLA-E/VL9 tetramer⁺ CD8⁺ T cells from CMV⁺ healthy donors D1, D5, D6, and D16 under CD94 blockade. Data are background-subtracted relative to unstimulated controls. Statistical significance is indicated as follows: ns, P > 0.05; *P ≤ 0.05; **P ≤ 0.01; ***P ≤ 0.001; ****P ≤ 0.0001*.

IL-15-associated programs have previously been linked to both protective MHC-E-restricted immunity^54^, and KIR⁺ T-cell function^55^. We therefore asked whether IL-15 could further enhance E-sensor responses to ‘increased self,’ given that E-sensors are restricted by HLA-E, express KIR, and display phenotypes consistent with heightened IL-15 sensitivity. PBMCs were pre-exposed to escalating concentrations of an IL-15-based superagonist (heterodimeric IL-15/IL-15Rα, hereinafter termed *het-IL-15*)^56^ prior to being exposed to HLA-E upregulation in the form of self-VL9/HLA-E tetrameric stimulation. Het-IL-15 drove steep dose-dependent amplification of activation and degranulation within the E-sensor compartment across all donors **(Fig. 3e,f)**.

To dissect the individual contributions of IL-15 signaling versus HLA-E upregulation in licensing E-sensor activation, we tested them separately and in combination. HLA-E upregulation or het-IL-15 alone induced modest E-sensor activity, whereas combined exposure drove strong synergy in all donors, with E-sensor response magnitudes exceeding additive effects by 3-fold on average, demonstrating potent cooperativity between increased self and inflammatory cytokine signaling **(Fig. 3g,h)**. Het-IL-15 also erased the hierarchy between high- and low-MFI VL9 variants, equalizing the level of E-sensor activation and degranulation, irrespective of HLA-E/VL9 tetramer staining intensity **(Fig. 3i)**. These findings support a threshold-based model in which disease-associated increases in HLA-E enable E-sensors to break tolerance and respond to ‘increased self’, while disease-associated inflammatory cues synergize with increased HLA-E to license potent E-sensor responses in the absence of foreign antigen.

### E-sensors mediate immunoregulatory activity through selective elimination of highly activated, proliferating immune cells

Immunoregulatory KIR^+^ CD8^+^ T cells have previously been associated with the prevention of autoimmune disease through homeostatic elimination of antigen-activated CD4^+^ T cells, although the mechanistic basis of recognition remains unknown^39^. Given that E-sensors also express inhibitory KIRs, which restrain autoreactivity against healthy self, and that CD4⁺ T-cell activation has previously been shown to induce HLA-E upregulation^43^, we hypothesized that E-sensors may mediate immunoregulatory killing of highly activated, proliferating CD4⁺ T cells through increased HLA-E expression. Consistent with prior reports^43^, we show that nonspecific CD4^+^ T cell activation increased HLA-E surface expression by approximately threefold relative to non-activated CD4⁺ T cells across all donors **(Fig. 4a)**. We next examined the immediate *ex vivo* elimination capacity of E-sensors using directly isolated, unmanipulated CD8⁺ T cells from healthy CMV-seropositive persons harboring expanded E-sensor populations (>1% of total CD8^+^ T cells) **(Fig. 4b)**. Addition of isolated E-sensor^+^ CD8^+^ effector cells resulted in selective and effector-dose-dependent elimination of activated CD4^+^ T cells **(Fig. 4c)**, which was highly reproducible across five independent experiments performed on separate days **(Fig. 4d).** Progressive depletion of activated CD4^+^ T cells over increasing CD8^+^ effector ratios was observed in all donors with E-sensor populations, reaching 57% elimination at the highest effector to target (E:T) ratio of 16:1, and was almost completely abrogated by HLA-E blockade **(Fig. 4e)**. Cumulative targeting (area under the elimination curve, AUC) and maximal elimination at E:T 16:1 were significantly reduced under HLA-E blockade (two-tailed t-tests, p < 0.0001), underscoring strict HLA-E-dependency of activated CD4^+^ T cell elimination **(Fig. 4f)**. We also depleted E-sensors using HLA-E/VL9 tetramers under CD94 blockade, coupled with bead-based or saporin-mediated depletion strategies **(Fig. 4g)**. E-sensor depletion resulted in significant reductions in elimination relative to mock-depleted controls **(Fig. 4h)**, whether measured by cumulative cytotoxicity (AUC) or maximal E: T ratios (p < 0.001), and the efficiency of E-sensor depletion was proportional to killing inhibition **(Fig. 4i,j)**, establishing an immunoregulatory role for E-sensors in controlling activated CD4^+^ T cell expansion.

**Figure 4.**
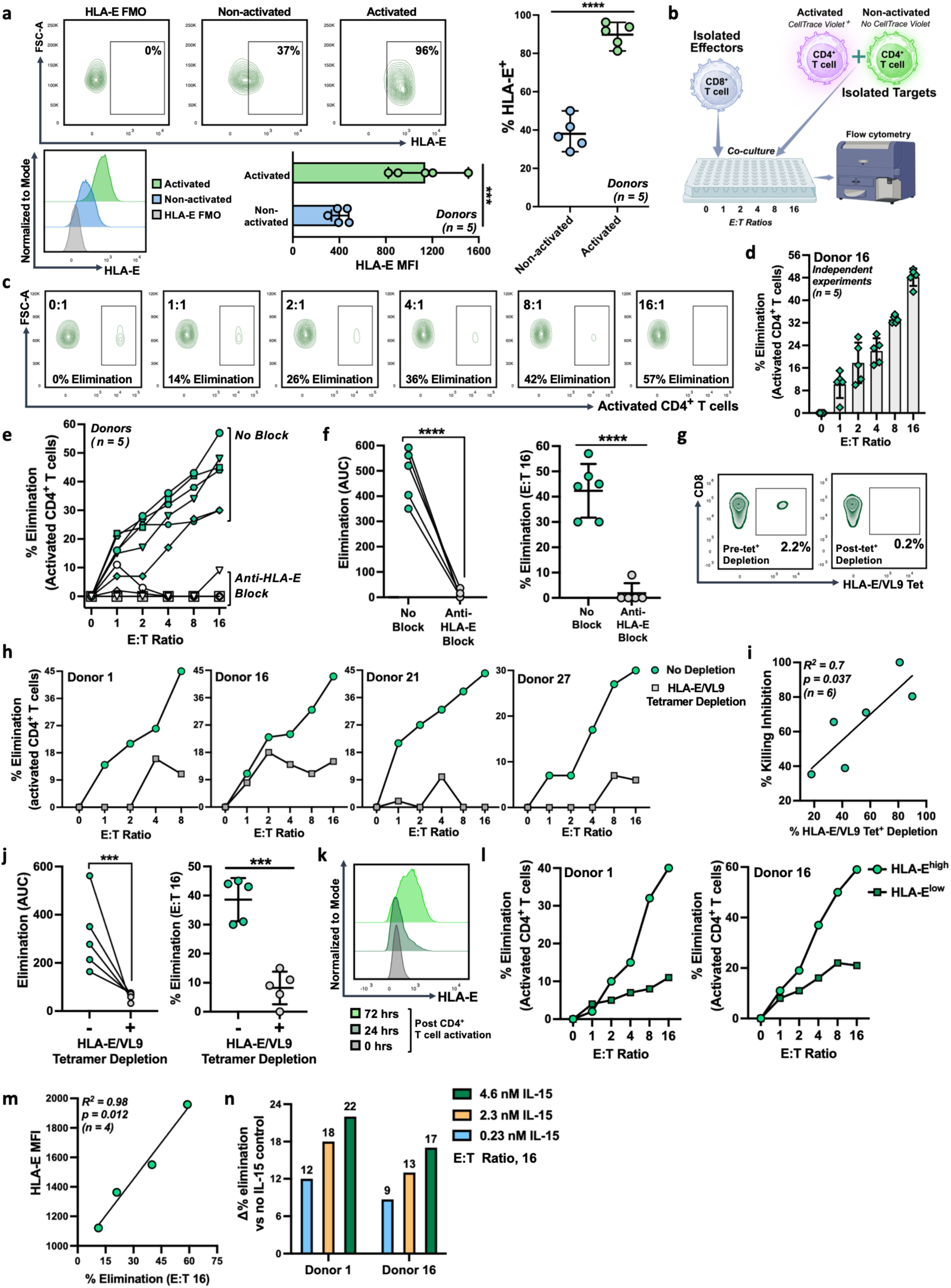
E-sensors mediate immunoregulatory activity via selective elimination of activated CD4^+^ T cells. **(a)** Top, representative HLA-E surface expression staining of live CD3⁺CD4⁺ T cells from a healthy donor. HLA-E-positive gates were defined using a fluorescence-minus-one (FMO) control (left). Non-activated CD4⁺ T cells (middle) and CD4⁺ T cells following 72 h anti-CD3/anti-CD28 activation (right) are shown. Bottom and right panel: representative histograms and quantification of HLA-E MFI and HLA-E⁺ frequency on activated versus non-activated CD4^+^ T cells from five healthy donors. **(b)** Schematic of the *ex vivo* immunoregulatory elimination assay. CD4⁺ T cells were activated for 72 h and mixed with freshly isolated non-activated CD4⁺ T cells. All CD4^+^ cells were labeled with CellTrace Far Red (CTFR), while activated CD4⁺ T cells were additionally labeled with CellTrace Violet (CTV). Mixed CD4^+^ target populations were co-cultured with purified CD8⁺ T-cell effectors across increasing effector-to-target (E:T) ratios. Following a 24 h co-culture, selective elimination of activated CD4⁺ T cells was quantified by flow cytometry as the frequency of CTV⁺ targets relative to non-activated targets. **(c)** Representative elimination assay plots across increasing E:T ratios gated on live single CD3^+^CD4^+^ T cell targets from Donor 16. Cells within the specified gate are CTV^+^ activated CD4^+^ T cells. Cells outside the gate are non-activated CD4^+^ T cells. **(d)** Percentage elimination of activated relative to non-activated CD4⁺ T cells from Donor 16 across increasing E:T ratios. Data represent five independent experiments performed on separate days. Symbols indicate individual experiments; bars show the mean ± SEM. **(e)** Elimination assay plot as in D, for five CMV⁺ donors with expanded E-sensor populations (>1% of CD8^+^ T cells). Green-filled symbols indicate no blockade; open symbols indicate anti-HLA-E blockade. Symbol shapes identify individual donors. **(f)** Effect of HLA-E blockade on elimination. Left, area under the elimination curve (AUC). Lines connect donor-matched measurements. Right, elimination at an E:T ratio of 16:1. Horizontal lines indicate medians; error bars indicate SD. Stars denote statistical significance by two-tailed paired t-tests. **(g)** Representative HLA-E/VL9 tetramer staining before and after tetramer-mediated E-sensor depletion. Frequencies are shown as a percentage of total CD8⁺ T cells. Plots are gated on single live CD3^+^CD8^+^ T cells. **(h)** Effect of E-sensor depletion on elimination of activated CD4⁺ T cells. Green symbols indicate elimination assays using total CD8⁺ T cells; grey symbols indicate CD8⁺ T cells following E-sensor depletion. Each plot represents an individual donor. **(i)** Correlation between E-sensor depletion efficiency and inhibition of elimination at an E:T ratio of 16:1. Each point represents an independent experiment. The line shows a simple linear regression (*R^2^ = 0.7, p = 0.037*). **(j)** As in F, except comparing conditions with and without E-sensor depletion. **(k)** Representative HLA-E expression on CD4⁺ T cells at 0, 24, and 72 h following non-specific activation. **(l)** Elimination of activated CD4⁺ T cells harvested 24 h (HLA-E^low^) or 72 h (HLA-E^high^) following activation. Each plot shows a separate donor. Percentage elimination was quantified relative to non-activated CD4⁺ T cells, as in D. **(m)** Correlation between HLA-E MFI and the corresponding percentage elimination at an E:T ratio of 16:1 for HLA-E^low^ and HLA-E^high^ CD4^+^ T cell targets from Donors 1 and 16. The line shows a simple linear regression (R² = 0.98; p = 0.012). **(n)** Effect of IL-15 superagonist on elimination of activated CD4⁺ T cells. Bars show the increase in percentage elimination relative to the no-IL-15-superagonist condition at an E:T ratio of 16:1 across increasing IL-15 superagonist concentrations. Values above bars indicate the increase in percentage elimination. *Unless otherwise specified, statistical significance was determined via two-tailed unpaired t-tests and is indicated as follows: ns, P > 0.05; *P ≤ 0.05; **P ≤ 0.01; ***P ≤ 0.001; ****P ≤ 0.0001*.

Notably, E-sensor killing scales with HLA-E expression levels on target cells. Nonspecific activation induced progressive increases in surface HLA-E expression on CD4^+^ T cells over time **(Fig. 4k)**, enabling comparison of E-sensor-mediated killing of target cells expressing progressively higher levels of HLA-E relative to the non-activated baseline. Activated CD4⁺ T cells from all post-activation time points were susceptible to E-sensor-mediated elimination relative to resting CD4^+^ T cells; however, elimination was consistently greater when target cells expressed higher levels of HLA-E **(Fig. 4l)**. Strikingly, target cell elimination exhibited a near-perfect linear relationship with HLA-E surface expression (R² = 0.98, P = 0.012), demonstrating that E-sensor cytotoxicity quantitatively scales with target cell HLA-E expression **(Fig. 4m)**.

Consistent with het-IL-15-induced E-sensor activation, het-IL-15 preconditioning increased elimination of activated CD4^+^ T cells in a dose-dependent manner across donors, enhancing cytotoxicity relative to untreated controls and demonstrating that disease-associated inflammatory cues can further amplify immunoregulatory elimination **(Fig. 4n)**. Together, these findings define an HLA-E-dependent, HLA-E-restricted immunoregulatory mechanism in which E-sensors selectively eliminate highly activated, proliferating immune cells in the absence of foreign antigen. Target-cell susceptibility closely tracked surface HLA-E expression, supporting a role for E-sensors in limiting excessive immune activation and restoring immune homeostasis.

### E-sensors detect pathogen-induced increases in HLA-E and eliminate infected cells

Given that pathogens frequently upregulate HLA-E^14,36,57^, we hypothesized that infected cells may also be intrinsically vulnerable to E-sensor targeting. We tested this using HIV as a model system and performed viral elimination assays using cells from four HIV-negative donors with expanded E-sensor populations. We infected CD4^+^ T cells with HIV *in vitro* and demonstrated progressive effector-dose-dependent depletion of virally infected cells relative to uninfected resting CD4^+^ T cells across all donors **(Fig. 5a-d)**. This immediate antiviral elimination was significantly reduced by HLA-E blockade and E-sensor depletion **(Fig. 5b-d)**.

**Figure 5.**
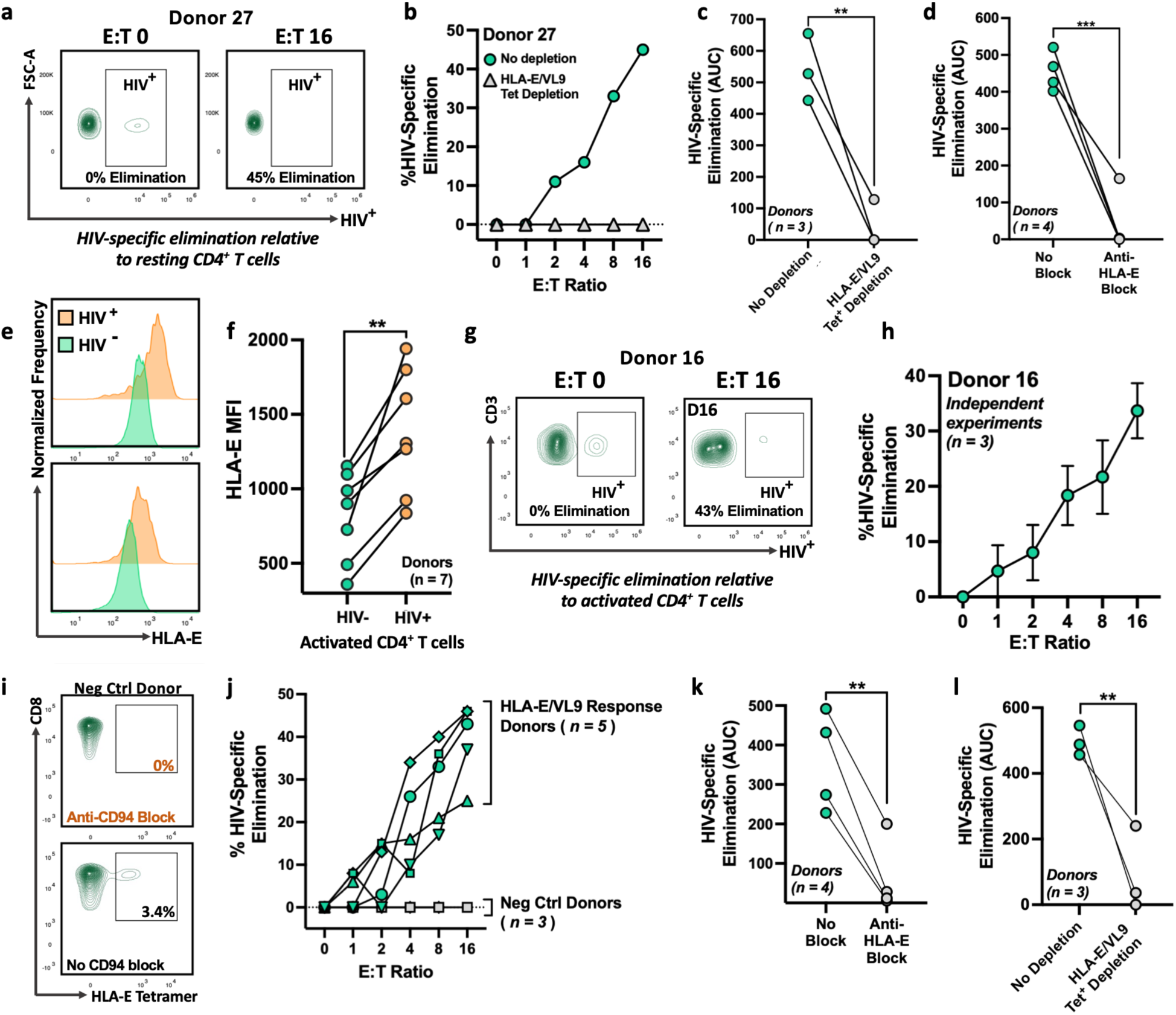
E-sensors from HIV-negative donors eliminate HIV-infected CD4⁺ T cells *ex vivo.* **(a)** Representative flow cytometry plots from a CMV seropositive HIV-negative donor (27) with a large (>1% of CD8^+^ T cells) E-sensor population, showing *ex vivo* elimination of HIV-infected CD4⁺ T cells. Pre-activated CD4⁺ T cells infected with VSV-G-pseudotyped HIV-1 NL4-3 ΔEnv GFP were mixed with resting CD4⁺ T cells and cultured in the absence (Target only, E:T = 0) or presence of E-sensors (E:T = 16:1). Plots are gated on live single CTFR⁺ target cells, with GFP identifying HIV-infected targets. **(b)** Representative donor (27) showing HIV-specific elimination relative to resting CD4⁺ T cells across increasing effector-to-target ratios, with or without E-sensor depletion. **(c,d)** Area under the HIV-specific elimination curve for assays performed as in a,b, following E-sensor depletion (c) or HLA-E blockade (d) in healthy CMV-seropositive donors with expanded E-sensor populations. Lines connect donor-matched measurements. **(e)** Histograms for two representative healthy donors showing HLA-E surface expression on isolated, pre-activated CD4⁺ T cells, 72 hours post HIV infection (NL4-3-ΔEnv-GFP virus pseudotyped with VSV-G) or mock infection. Gated on live singlet CD3^+^ CD8^-^ T cells. **(f)** HLA-E MFI on isolated pre-activated CD4⁺ T cells following HIV infection or mock infection in seven healthy donors. Lines connect donor-matched samples; statistical significance from two-tailed unpaired t-tests is indicated. **(g)** Flow cytometry plots from an HIV elimination assay on a representative donor (D16) with an expanded (>1%) E-sensor population. Pre-activated HIV-infected and pre-activated mock-infected CD4⁺ T cells were labeled with cell-specific dyes, mixed, and co-cultured with isolated CD8⁺ T-cell effectors. Plots are gated on live single CD3⁺CTFR⁺ target cells and show HIV-infection rates (ascertained by HIV-associated GFP expression) at E:T ratios of 0:1 and 16:1. **(h)** Selective elimination of pre-activated HIV-infected CD4⁺ T cells relative to pre-activated mock-infected CD4⁺ T cells across increasing E:T ratios. Data represent three independent experiments performed on separate days for a representative donor (D16). Symbols represent the mean from three individual experiments, and error bars denote SEM. **(i)** Representative HLA-E/VL9 tetramer staining from a healthy negative control donor lacking a detectable E-sensor population, with (top) or without (bottom) CD94 blockade. Plots are gated on live singlet CD3⁺CD8⁺ T cells. Gates indicate HLA-E/VL9 tetramer-positive cells, with frequencies shown as a percentage of total CD8⁺ T cells. **(j)** HIV-specific elimination relative to pre-activated mock-infected CD4^+^ T cells, as in (h). Green symbols represent donors with highly expanded E-sensor populations (n = 5), whereas grey symbols represent healthy control donors lacking detectable E-sensor populations (n = 3). **(k, l)** Area under the HIV-specific elimination curve for assays performed as in (g,h,j), following anti-HLA-E blockade (k) or E-sensor depletion (l) in CMV seropositive donors with large (>1%) E-sensor populations. Lines connect donor-matched measurements. *Statistical significance in this figure was determined using two-tailed unpaired t-tests and is indicated as follows: ns, P > 0.05; *P ≤ 0.05; **P ≤ 0.01; ***P ≤ 0.001; ****P ≤ 0.0001*.

We next asked whether the preferential elimination of HIV-infected targets would persist when quantified relative to non-infected CD4⁺ T cells that had been activated identically to their HIV-infected counterparts. Significantly higher HLA-E expression was observed on activated HIV-infected CD4^+^ cells than on mock-infected activated controls across all donors, suggesting that infection may further enhance susceptibility to E-sensor targeting **(Fig. 5e,f)**. *Ex vivo* elimination assays were performed to quantify the relative depletion of HIV-infected versus HIV-negative activated CD4⁺ T cells. Progressive and reproducible, effector-dose-dependent depletion of HIV-infected cells relative to uninfected counterparts was observed **(Fig. 5g,h)**. This phenotype was observed across multiple HIV-negative HCMV seropositive donors with expanded E-sensor populations but was absent in donors lacking detectable E-sensor responses **(Fig. 5i,j)**, and was significantly reduced by HLA-E blockade **(Fig. 5k)** and by E-sensor depletion **(Fig. 5l)**. Together, these findings demonstrate that E-sensors from HIV-negative individuals can directly and preferentially eliminate HIV-infected CD4⁺ T cells *ex vivo* in an HLA-E-dependent manner despite no prior HIV antigen exposure, suggesting a potential role for these cells in contributing to spontaneous HIV immune control. More broadly, these results reveal an innate-like capacity of E-sensors to mediate immunosurveillance of pathogen-induced increases in HLA-E expression in the absence of prior pathogen-specific priming.

### E-sensors are enriched in HIV elite controllers and define a distinct mechanism of HIV immune control

Elite controllers, comprising <0.5% of HIV-infected individuals^58^, spontaneously suppress viral replication in the absence of antiretroviral therapy, yet the mechanisms underlying this rare, heterogeneous phenotype remain incompletely understood. Although classical HLA class I-restricted HIV-specific CD8⁺ T-cell responses are strongly associated with viral control, known genetic determinants account for only ∼20% of the elite-control phenotype, and no single mechanism explains control across all individuals^59,60^. Given the rapid dissemination of HIV following infection, we reasoned that pre-existing, effector-ready E-sensors capable of immediate antiviral activity could provide an additional, kinetically advantaged component of HIV immune control that does not require HIV-specific T-cell priming, clonal expansion, or effector differentiation.

We screened a cohort of HIV elite controllers for E-sensor responses. Staining with self-VL9/HLA-E tetramers not only confirmed that E-sensors in HIV elite controllers were self-reactive **(Fig. 6a)**, mirroring the self-VL9 reactivity observed in HCMV-seropositive HIV-negative donors, but also demonstrated their longitudinal persistence, with elevated frequencies maintained across all time points spanning more than 13 years **(Fig. 6b)**. Distinguishable MFI-separated E-sensor populations were detected in 13 of 24 (54%) HIV elite controllers **(Fig. 6c)** and ranged in frequency from small but clearly distinguishable populations (∼0.2% of CD8⁺ T cells) to exceptionally large expansions approaching 12% of the CD8⁺ T-cell repertoire.

**Figure 6.**
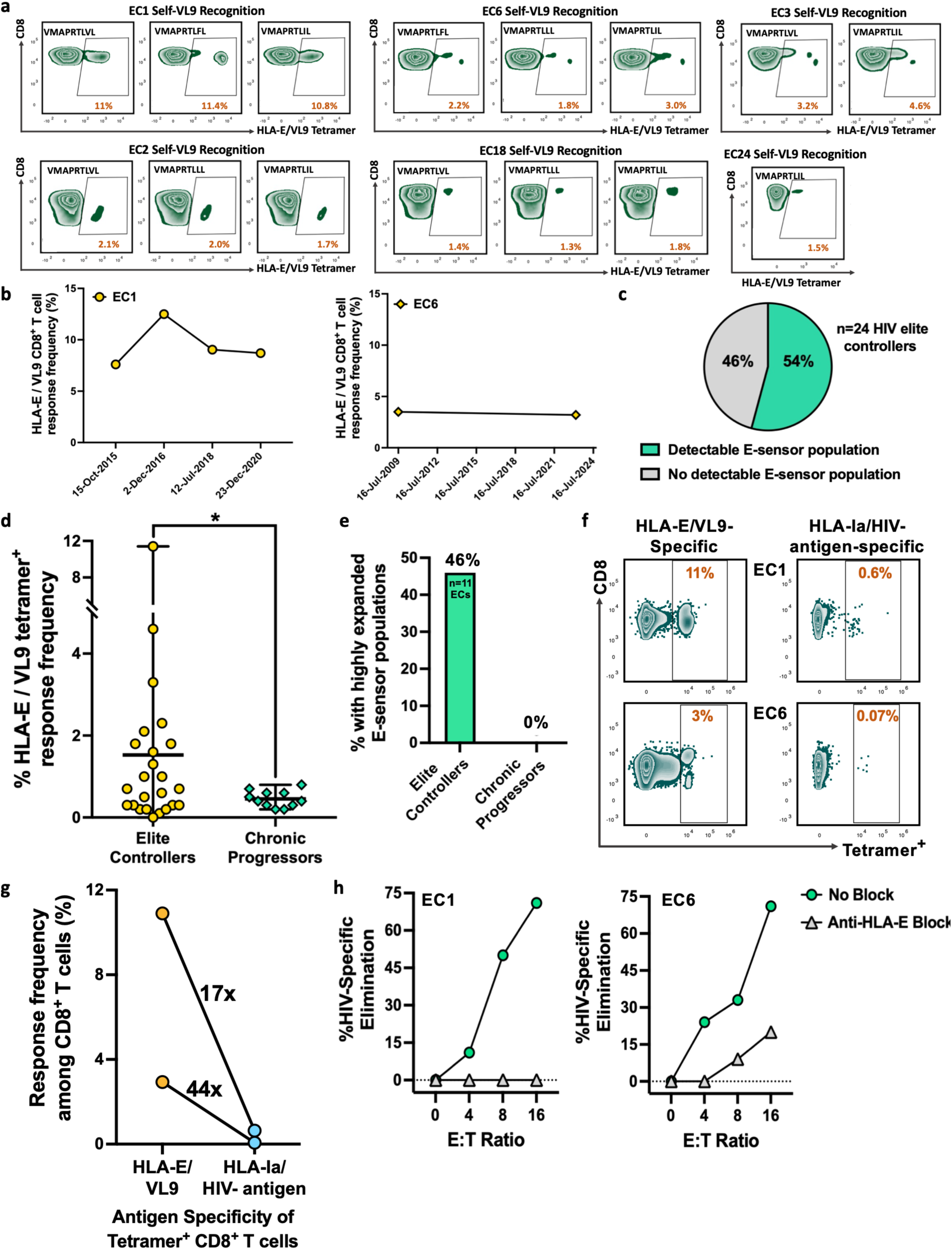
E-sensors are enriched in HIV elite controllers and mediate HLA-E-dependent elimination of HIV-infected cells. **(a)** Representative HLA-E tetramer staining with donor-matched self-VL9 peptides under CD94 blockade from six HIV elite controllers with distinct E-sensor populations. Plots are gated on live single CD3⁺CD8⁺ T cells, and frequencies indicate tetramer^+^ cells as a percentage of total CD8⁺ T cells. **(b)** Longitudinal frequencies of expanded HLA-E/VL9 tetramer^+^ CD8⁺ T cell responses in two HIV elite controllers. Tetramer staining was performed as in B. Symbols represent individual time points. **(c)** Proportion of HIV elite controllers (n = 24) with a distinguishable MFI-separated HLA-E/VL9 E-sensor population. **(d)** Frequency of HLA-E/VL9 tetramer^+^ E-sensors in HIV^+^ elite controllers (n=24) and chronic progressors (n=11). Symbols represent individual donors; horizontal lines indicate the median, and error bars denote the range. Statistical significance was determined by a two-tailed unpaired t-test. **(e)** Percentage of HIV elite controllers and chronic progressors with highly expanded E-sensor populations comprising >1% of CD8^+^ T cells. **(f)** Direct *ex vivo* tetramer staining for elite controller 1 (EC1) and 6 (EC6). E-sensor populations are shown (left), alongside conventional HIV-antigen-specific responses restricted by classical HLA class I (right). Conventional HIV-specific responses were restricted by HLA-A11:01 (AK11) or HLA-B57:01 (KF11). Plots are gated on live single CD3⁺CD8⁺ T cells, and frequencies indicate tetramer^+^ cells as a percentage of total CD8⁺ T cells. **(g)** Frequencies of HLA-E/VL9 E-sensor and conventional HIV-specific tetramer-positive CD8⁺ T-cell responses in HIV elite controllers as in (f). Lines connect paired measurements from the same donor. Fold differences between paired responses are indicated. **(h)** As in Fig. 5g-l, HIV-specific elimination of infected CD4^+^ T cells relative to mock-infected counterparts by CD8⁺ T cells from HIV^+^ elite controllers in the presence or absence of anti-HLA-E blockade. *Statistical significance is indicated as follows: ns, P > 0.05; *P ≤ 0.05; **P ≤ 0.01; ***P ≤ 0.001; ****P ≤ 0.0001*.

Notably, E-sensor responses were enriched in HIV elite controllers relative to persons who do not naturally control HIV infection and require antiretroviral therapy to suppress progressive infection (two-tailed t-test, P = 0.0392) **(Fig. 6d)**. Furthermore, highly expanded E-sensor populations comprising 1% or more of CD8⁺ T cells were present in 46% of elite controllers but absent in persons with antiretrovirally suppressed progressive infection **(Fig. 6e)**. To place the magnitude of elite controller E-sensor responses into context, we directly compared HLA-E/VL9 tetramer^+^ E-sensor frequencies with conventional HIV-antigen-specific CD8⁺ T-cell responses in the same elite controllers using classical HLA-A*11:01/AK11 and HLA-B*57:01/KF11 tetramers. Strikingly, E-sensor populations were 17- and 44-fold larger than the corresponding HIV-antigen-specific CD8⁺ T-cell responses **(Fig. 6f,g)**. In both controllers, conventional HIV-specific responses were readily distinguishable but comprised less than 1% of CD8⁺ T cells, with one accounting for less than 0.1%.

We next investigated whether these expanded E-sensor populations mediated immediate antiviral activity in HIV elite controllers. HIV elimination assays using elite controllers with expanded E-sensor populations revealed HLA-E-dependent elimination of HIV-infected CD4⁺ T cells **(Fig. 6h)**. In one elite controller, whose E-sensor population comprised more than 11% of the total CD8⁺ T cell repertoire, HLA-E blockade completely abrogated elimination, indicating that immediate cytotoxicity was entirely attributable to HLA-E-mediated responses. In a second elite controller, HLA-E blockade partially reduced elimination, suggesting the presence of additional antiviral CD8⁺ T cell populations capable of mediating immediate *ex vivo* effector function. Together, these findings identify E-sensors as a durable, frequently expanded, and functionally poised CD8⁺ T cell population in a model chronic viral infection and implicate HLA-E-restricted immunity as a previously unrecognized innate-like axis of HIV immune control.

## DISCUSSION

Here, we establish a new pathway of human CD8^+^ T cell immunosurveillance that couples antiviral defense and immunoregulation through recognition of disease-associated increases in the constitutively expressed non-classical HLA class I molecule, HLA-E. This represents a previously unrecognized mode of human T cell immunosurveillance, in which TCR-mediated quantitative sensing of a constitutively expressed self-ligand is balanced by inhibitory KIR signaling, enabling detection of pathological ‘increased self’ while maintaining tolerance to healthy self. This mode of recognition contrasts with conventional CD8^+^ T cell and NK cell immunity, which typically recognize foreign antigen presented by classical HLA-A, -B, or -C, or ‘missing self’ through loss of HLA class I expression, respectively. It also differs from surveillance pathways that recognize ‘induced self’ following stress or transformation: E-sensors instead detect quantitative increases in a self-ligand that is constitutively expressed and required for immune homeostasis.

E-sensors co-express both activating and inhibitory receptors that recognize distinct HLA class I molecules: activating TCRs that recognize self-VL9/HLA-E, and inhibitory KIRs that bind HLA-C^37^. Under our model, healthy cells express lower levels of HLA-E, which generate limited TCR-mediated activating signals, whereas higher levels of HLA-C engage inhibitory KIRs, enforcing self-tolerance. In contrast, altered and disease-associated states frequently upregulate HLA-E^14,33,34,61,62^, thereby enhancing activating signals through self-VL9/HLA-E-reactive TCRs and shifting the balance from self-tolerance toward immune surveillance, enabling the elimination of pathological cells. This effect may be further amplified by disease-associated inflammatory cues, as we demonstrate for IL-15, providing an additional layer of contextual regulation that may preferentially potentiate E-sensor activity in disease. E-sensor activation may also be favored in settings where KIR-mediated inhibitory signaling is reduced, such as pathogen-mediated HLA-C downregulation^44^ or stress-induced remodeling of the HLA-C peptidome^45^, which can diminish inhibitory KIR engagement. Together, our findings support a model in which E-sensors detect ‘increased self’ through quantitative sensing of self-VL9/HLA-E, with the resulting activating signal balanced against inhibitory KIR signaling and modulated by inflammatory cues to calibrate the transition from self-tolerance to immune surveillance **(Fig. 7)**.

**Figure 7.**
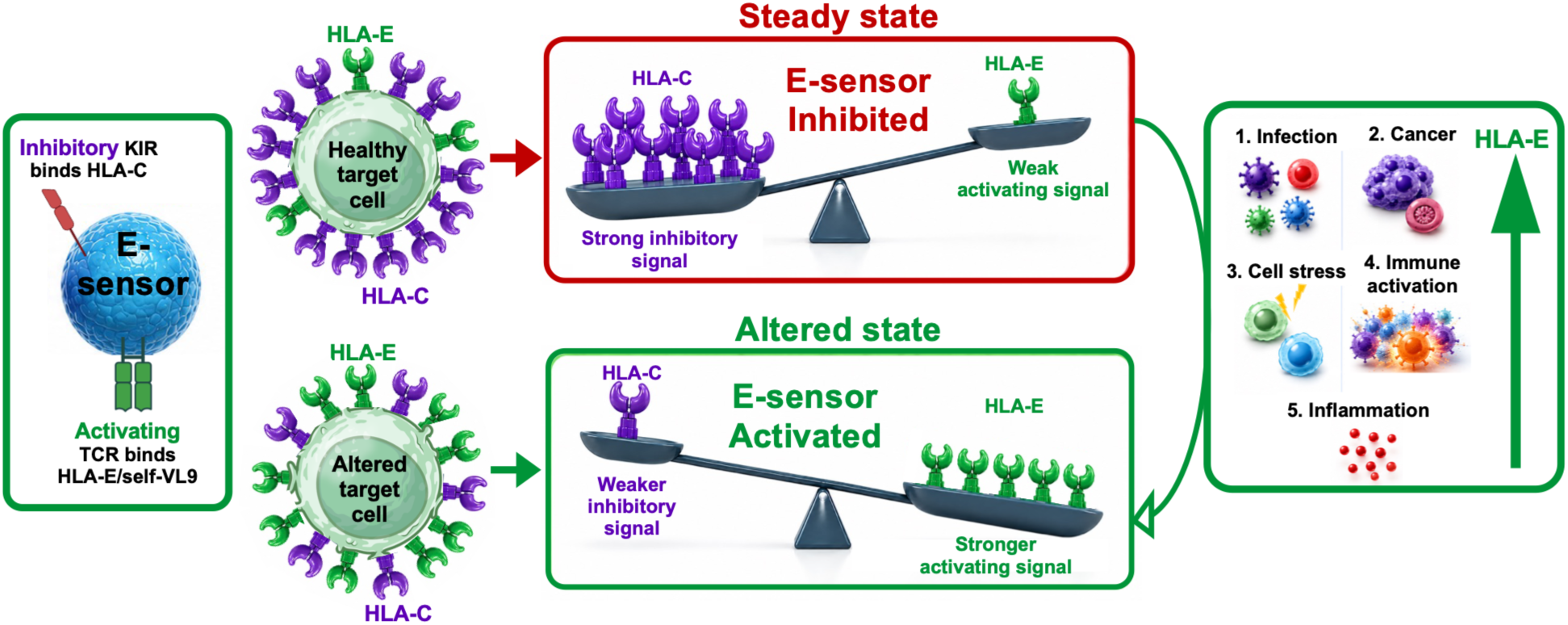
A new pathway of HLA-E-restricted CD8^+^ T cell-mediated immunosurveillance and immunoregulation. E-sensors are HLA-E-restricted VL9-specific CD8^+^ T cells that co-express activating TCR that recognize self-VL9/HLA-E and inhibitory KIR receptors that recognize HLA-C, thereby integrating opposing signals from distinct MHC class I ligands. Under homeostatic conditions, healthy cells express lower HLA-E relative to HLA-C, resulting in limited self-VL9/HLA-E—TCR-mediated activating signal and dominant HLA-C—KIR-mediated inhibition, thereby promoting self-tolerance. In contrast, altered and disease-associated cellular states—including infection, malignancy, cellular stress, inflammation, and immune-cell activation—are often associated with increased HLA-E surface expression. Under our model, such pathological increases in HLA-E abundance enhance engagement of the activating HLA-E/VL9-specific TCR, shifting the balance from self-tolerance toward immunosurveillance, and enabling the elimination of pathological cells. This effect may be further amplified in settings where inhibitory KIR signaling is reduced, such as pathogen-mediated HLA-C downregulation or stress-induced alterations in the HLA-C peptidome that weaken inhibitory KIR engagement. Together, these findings support a model in which E-sensors detect ‘increased self’, enabling immunoregulatory and antiviral immunosurveillance while maintaining tolerance toward healthy self.

Although KIR^+^ HLA-E/VL9-specific CD8⁺ T-cell responses were previously characterized as HCMV-specific responses^37,38,63^, we demonstrate that these cells mediate functions beyond CMV immune control. In contrast to well-established CD4⁺ regulatory T cells, regulatory CD8⁺ T cells remain comparatively poorly characterized, with a distinct population of KIR⁺ regulatory CD8⁺ T cells only recently robustly defined^39^. However, the antigen specificity of these cells and the molecular basis of their target cell recognition remained unknown. Our findings build on these prior observations by establishing a mechanistically resolved pathway of immunosurveillance and immunoregulation mediated by KIR^+^ HLA-E-restricted self-VL9-reactive CD8^+^ T cells.

The quantitative relationship between activation-induced HLA-E expression and E-sensor-mediated elimination suggests a graded homeostatic negative-feedback mechanism in which the most highly activated immune cells become preferentially susceptible to regulatory cytotoxicity. The amplification of E-sensor activity by IL-15 suggests that the strength of this regulatory response may also be coupled to inflammatory context, potentially enabling regulation to scale with the magnitude of immune activation. Such a mechanism could help prevent excessive immune activation and promote the restoration of immune homeostasis. This model is consistent with previous *in vivo* evidence that loss of regulatory KIR⁺ CD8⁺ T cells permits pathological expansion of activated CD4⁺ T cells and can otherwise lead to autoimmune disease^39^. However, in contrast to the well-established identity and suppressive mechanisms of CD4⁺ regulatory T cells, the existence and mechanistic basis of regulatory CD8⁺ T cells have remained elusive for decades, particularly in humans. E-sensors represent a mechanistically defined population within this broader KIR⁺ regulatory CD8⁺ T-cell compartment, providing a molecular basis for discrimination of activated targets through quantitative sensing of HLA-E/VL9. By linking antigen specificity to regulatory cytotoxic function, our findings identify a mechanistically resolved pathway of human CD8⁺ T cell immunoregulation.

We also show that E-sensors detect pathogen-induced increases in HLA-E expression, using HIV as a model system. Our findings provide a mechanistic framework that may help reconcile the identification of diverse pathogen-derived ligands proposed to mediate MHC-E-restricted CD8⁺ T-cell responses, many of which lack canonical peptide sequence motifs or evidence of stable presentation under physiological conditions^9,49,64,65^. Our findings therefore expand the mechanistic framework for HLA-E-restricted immunity by demonstrating that pathogen-induced increases in the canonical self-VL9/HLA-E can themselves provide a signal for antiviral T-cell recognition.

Although HLA-E-restricted HIV-specific responses have been reported, the proposed epitopes exhibit limited HLA-E stability or evidence of endogenous presentation^7,26,28,64,66,67^. Our data offer an alternative mechanistic paradigm: HIV preferentially infects activated CD4⁺ T cells, that express increased levels of HLA-E, and infection further amplifies HLA-E surface expression, further increasing susceptibility to E-sensor-mediated elimination. Moreover, because activated CD4⁺ T cells are the principal targets of HIV infection^68,69^, E-sensors may indirectly contribute to HIV immune control by restricting the pool of HIV-susceptible targets through control of activated CD4^+^ T cell expansion, thereby complementing their direct antiviral role. The enrichment of E-sensors in HIV elite controllers, together with their capacity to eliminate HIV-infected targets *ex vivo*— even in HIV-naïve individuals—supports a kinetically advantaged model of pre-positioned immune surveillance capable of acting immediately without prior HIV antigen exposure before conventional pathogen-specific responses are fully established.

This pre-positioned effector-ready state is reminiscent of prophylactic HIV vaccine models in which unprecedented immune clearance is mediated by MHC-E-restricted CD8^+^ T cell responses^19,25^. Intriguingly, induction of these vaccine responses—and the associated protection—is strictly dependent on a VL9-mimetic peptide encoded by the CMV-based vaccine vector; deletion of vaccine-encoded VL9 abrogates both MHC-E-restricted responses and vaccine protection^7070^. Yet, the mechanistic basis for this striking dependence on VL9 remains incompletely understood. Moreover, vaccine-mediated protection is associated not only with the induction of MHC-E-restricted CD8⁺ T cell responses but also with a sustained innate immune program involving IL-15-associated genes, which predicts vaccine efficacy^71^. In parallel, and consistent with prior reports showing that IL-15 lowers the activation threshold of KIR^+^ T cells^55,72^, we find that IL-15 further potentiates E-sensor responses to ‘increased self’. Thus, several features associated with protective vaccine immunity converge with E-sensor biology, including CMV association, MHC-E restriction^19^, terminally differentiated effector phenotypes^21^, IL-15-driven programs^71^, and VL9-linked pathways^70^. These parallels raise the possibility that related principles of MHC-E-restricted immune surveillance may operate in protective vaccine immunity, an important question for future investigation as this vaccine platform advances in human clinical trials.

The convergence of diverse biological processes—including infection^14,36,57^, inflammation^42^, immune activation^43^, cellular stress^33^, immune aging^73^, and cancer^33,62,74,75^—on HLA-E upregulation suggests that HLA-E functions as a shared molecular indicator of altered cellular state. Consistent with this, genome-wide *in vivo* screens identified the murine HLA-E ortholog, Qa-1, as one of the few shared determinants of resistance to immune-mediated tumor control among otherwise largely tumor-specific mechanisms^76^. Because E-sensors recognize this conserved host-derived signal rather than a disease-specific antigen, their functional relevance may extend to disease settings beyond those examined here. Disease-associated HLA-E upregulation typically drives immune inhibition via the CD94/NKG2A axis^1,33^, and in many settings, has been linked to poor clinical outcomes^33,62,77–80^. The clinical relevance of this inhibitory axis is underscored by therapeutic targeting of HLA-E-mediated inhibition in cancer through NKG2A blockade with the checkpoint antibody monalizumab^81^. Our model suggests that E-sensors are uniquely positioned to instead exploit pathological HLA-E upregulation, transforming this conserved mechanism of immune evasion and inhibition into a substrate for immune detection.

Conventional T cell-based vaccines and therapeutics rely on disease-specific antigens and are further constrained by the extensive polymorphism of classical HLA molecules. In contrast, E-sensors recognize a conserved marker of altered cellular state in a disease-agnostic manner, and HLA-E has only two major alleles in the human population, differing by a single amino acid outside the peptide-binding groove^82,83^. These features raise the possibility that E-sensor-directed immune interventions could achieve broader applicability across diseases and genetically diverse populations. More broadly, our findings define a form of immunosurveillance in which quantitative increases in constitutively expressed self, rather than foreign or inducible antigen or loss of self, provide the signal for cytotoxic recognition. This framework expands the functional scope of HLA-E-restricted immunity and provides a rationale for developing HLA-E-based interventions for diseases characterized by pathological HLA-E upregulation.

### Limitations

This study has several limitations that define clear directions for future work. First, while KIR⁺ CD8⁺ T cells have established immunoregulatory roles *in vivo*, the extent to which this HLA-E/VL9-reactive population specifically contributes to immune regulation *in vivo* will require expanded studies. Similarly, although E-sensors eliminated HIV-infected CD4⁺ T cells *ex vivo* in an HLA-E-dependent manner, establishing their contribution to pathogen control *in vivo* will be an important next step. Additionally, because E-sensor recognition depends on self-VL9/HLA-E upregulation rather than a disease-specific antigen, this mechanism may extend to other HLA-E-upregulating conditions, including other infections, cancer, autoimmune and inflammatory diseases, and immune aging; testing this generalizability is beyond the scope of the current study and an important direction for future work. Finally, whether similar responses contribute to RhCMV/SIV vaccine-elicited protection or to other previously described HLA-E-restricted responses is a particularly compelling question raised by our findings and warrants future investigation.

## MATERIALS AND METHODS

### Prokaryotic protein expression and purification

Recombinant human and rhesus β2-microglobulin (β2M), HLA-E\**01:03,* Mamu-E*02:04, and HLA-A*\**02:01 heavy chain constructs were cloned into pET-29b(+) expression vectors and expressed in *E. coli* BL21(DE3)pLysS (Thermo Fisher). Chemically competent cells were heat-shocked, recovered in SOC medium, and plated or directly introduced into overnight starter cultures. Overnight starters were grown in Luria broth (LB) (Research Products International) supplemented with kanamycin (100 µg/mL) at 37°C with agitation (80– 100 rpm). 10 mL was inoculated into 1 L of LB + kanamycin (100 µg/mL), and cultures were grown at 37°C (200 rpm) to an OD600 ≈ 0.5. Protein expression was induced with IPTG (0.5 mM final concentration) for 4 h at 37°C. Bacterial cells were harvested by centrifugation (8,000 rpm, 12 min, 4°C), and pellets were resuspended in lysis buffer (50 mM Tris-HCl, 0.7 M sucrose, 1 mM EDTA) supplemented with 10 mM DTT. Lysozyme (3.5 µM), DNase I (64.5 µM), and MgCl₂ (0.2 mM) were added, and suspensions were stirred for 10 min. EDTA was then increased to 10 mM, and inclusion bodies were pelleted by centrifugation (8,000 rpm, 20 min). Inclusion bodies were washed three times using a Triton-containing buffer (50 mM Tris-HCl, 0.5% Triton X-100, 100 mM NaCl, 1 mM EDTA, 2 mM DTT), followed by two washes with detergent-free buffer (50 mM Tris-HCl, 100 mM NaCl, 2.5 mM EDTA, 0.25 mM DTT). Pellets were solubilized in 8 M urea overnight at 4°C. Insoluble material was removed by centrifugation (19,000 rpm, 20–30 min, 4°C). Protein concentrations were determined by absorbance at 280 nm using extinction coefficients. Proteins were aliquoted, snap-frozen in liquid nitrogen, and stored at −80°C.

### Peptides

Synthetic peptides were produced using Fmoc (9-fluorenylmethoxycarbonyl) chemistry to >85% purity by GenScript and provided as lyophilized powder. Peptides were reconstituted in DMSO at 200 mM and stored at −80°C.

### MHC class I protein refolding

Refolding of peptide-HLA complexes (HLA-E*01:03, Mamu-E*02:04, and HLA-A*02:01) was performed at 4°C, over 72 hrs in refolding buffer containing 100 mM Tris-HCl, 400 mM L-arginine, 2 mM EDTA, 5 mM reduced glutathione, and 0.5 mM oxidized glutathione. Denatured species-matched β2M was first added by serial dilution to a final concentration of 2 µM, followed by peptide (added after ∼30 min incubation at 10-60 µM), and then HLA heavy chain (to a final concentration of 1 µM) using the same controlled dilution approach to minimize aggregation. After a 72 h incubation period, solutions were filtered (1.0 µM cellular nitrate membranes) and concentrated using tangential flow filtration (10 kDa MWCO, VivaFlow 50R system, Sartorius), followed by centrifugal concentration (10 kDa MWCO, VivaSpin Turbo Ultrafiltration). Refolded complexes were purified by size exclusion chromatography (Superdex S200 Increase 10/300 GL, Cytiva) in 20 mM Tris-HCl and 100 mM NaCl. Fractions corresponding to correctly folded peptide-HLA-β2M complexes were collected. Protein concentrations were determined by absorbance at 280 nm using extinction coefficients prior to biotinylation.

### MHC-E thermostability assays

Pre-refolded peptide-free HLA-E*01:03/β2-microglobulin and Mamu-E*02:04/β2-microglobulin complexes were incubated at room temperature with test peptides at concentrations ranging from 9–75 µM, as previously described^26^. Reactions were prepared in buffer containing 400 mM L-arginine monohydrochloride (Sigma-Aldrich), 5 mM reduced glutathione (Sigma-Aldrich), 0.5 mM oxidized glutathione (Sigma-Aldrich), 2 mM EDTA (Invitrogen), and 100 mM Tris-HCl (pH 8.0; Invitrogen). All conditions were performed in triplicate. Thermal stability was assessed by differential scanning fluorimetry using a Prometheus NT.Plex instrument (NanoTemper). Samples were loaded into nanoDSF-grade standard capillaries (NanoTemper), and excitation intensity was adjusted to 30–40% to maintain fluorescence signals between 5,000 and 15,000 arbitrary fluorescence units at both 330 and 350 nm. Samples were heated from 15°C to 95°C at a rate of 1°C/min. Melting temperatures (T_m_) were determined from the inflection point of the 350/330 nm fluorescence ratio using PR.ThermControl software (NanoTemper).

### Immunopeptidomics

HLA class I-deficient K562 cells stably overexpressing HLA-E*01:03 (kindly provided by the Birnbaum laboratory) were infected with VSV-G pseudotyped HIV (generated as described below) or mock-infected in multiple independent experiments. Successful infection was confirmed by GFP expression. Between 5×10^8^ and 8×10^8^ cells were harvested per condition, washed with PBS, lysed in 1% CHAPS-containing lysis buffer supplemented with protease inhibitors, and clarified by centrifugation. Lysates were subjected to immunoprecipitation using the anti-HLA-E monoclonal antibody 3D12 coupled to Protein A Sepharose beads. Following extensive washing, HLA-E-bound peptides were eluted with 10% acetic acid, separated from HLA-E heavy chain and β2-microglobulin using 10-kDa molecular weight cut-off filters (Pall), snap-frozen in liquid nitrogen, lyophilized, and stored at −80 °C prior to mass spectrometric analysis.

### DDA mass spectrometry analyses

HLA-E peptide samples were resuspended in 5 µL 0.1% formic acid. For all MS analyses, samples were analyzed using an Orbitrap Exploris 480 mass spectrometer (Thermo Fisher Scientific) coupled with an UltiMate 3000 RSLC Nano LC system (Dionex), Nanospray Flex ion source (Thermo Fisher Scientific), and column oven heater (Sonation). The MHC peptide sample was loaded onto a fused silica capillary chromatography column with an integrated electrospray tip (∼1 µm orifice) prepared and packed in-house with 10 cm of 1.9 µm C18 beads (ReproSil-Pur). Standard mass spectrometry parameters were as follows: spray voltage 2.0 kV, no sheath or auxiliary gas flow, ion transfer tube temperature 280 °C, RF lens 40%. The Orbitrap Exploris 480 mass spectrometer was operated in data dependent acquisition (DDA) mode. Peptides were eluted using the following gradient using 0.1% formic acid as buffer A and 70% Acetonitrile, 0.1% formic acid as buffer B: 6–25% buffer B over 75 min, 25–45% over 5 min, 45–97% over 5 min, hold for 1 min, and 97% to 3% over 2 min. Full scan mass spectra (350–1800 m/z, 60,000 resolution) were detected in the orbitrap analyzer after accumulation of 3x106 ions (i.e., normalized automatic gain control [AGC] target of 300%) or 25 ms. For every full scan, MS/MS scans were collected during a 3 s cycle time. Ions were isolated (0.4 m/z isolation width) using the standard (100%) AGC target, and automatic determination of maximum ion injection time (IT), fragmented by HCD with 30% collision energy (CE), and scanned at a resolution of 120,000. Charge states <2 and >4 were excluded, and precursors were excluded from selection for 30 s if fragmented n=2 times within a 20 s window.

### SureQuant mass spectrometry analyses

100 fmol of each synthetic SIL trigger peptide was spiked into each sample. Standard MS parameters and LC gradient were as described above (see DDA MS analyses), with the following modifications: MS1 scans were collected using a scan range of m/z 380-1200, a resolution of 120,000, an AGC target of 100%, and automatic determination of IT. Trigger peptide MS2 scans were collected at a resolution of 15,000, 27% CE, 100% AGC target, and automatic determination of IT. Target biological peptide MS2 scans were collected at a resolution of 240,000, 27% CE, 1000% AGC target, and 1000 ms max IT to maximize sensitivity. An LC flow rate of 300 nL/min was used. The custom SureQuant acquisition template available in the Thermo Orbitrap Exploris Series 2.0 method editor was used to build an analysis method as previously described^46,47^, targeting putative HIV-derived HLA-E-restricted peptides. SIL peptide precursor ions were matched with a mass accuracy tolerance of 10 ppm, and a threshold of n=3 out the top 6 product ions was used for pseudo-spectral matching with a mass accuracy tolerance of 20 ppm.

### MHC class I biotinylation and tetramer generation

Purified and refolded peptide-HLA class I monomers were enzymatically biotinylated using a commercial BirA-based biotinylation kit (Avidity), according to manufacturer instructions. Briefly, purified monomer was incubated with BirA enzyme, biotin, ATP, and magnesium at 4°C for ∼12 h, followed by removal of excess biotin by size exclusion chromatography (Superdex S200 Increase 10/300 GL, Cytiva). Aliquots corresponding to biotinylated monomer were snap-frozen and stored at −80°C. Tetramers were generated immediately prior to use by incubating 2.5µM of biotinylated monomer with fluorophore-conjugated streptavidin (BioLegend) at a final working concentration of ∼500 nM tetramer, ensuring excess monomer to saturate all four streptavidin binding sites.

### Human samples

Cryopreserved PBMCs from healthy CMV-seropositive donors were obtained from STEMCELL Technologies or isolated in-house from leukapheresis products. Leukapheresis products (leukopaks) were processed immediately upon receipt and peripheral blood mononuclear cells (PBMCs) were isolated using a magnetic-based direct isolation kit according to the manufacturer’s instructions (StemCell Technologies). Briefly, leukopak samples were diluted in phosphate-buffered saline (PBS) supplemented with 2% fetal bovine serum (FBS) and incubated with isolation cocktail and magnetic particles to deplete unwanted cell populations. Samples were then placed in a magnetic separation device (StemCell Technologies), and the enriched PBMC fraction was collected. Isolated PBMCs were washed, counted, and resuspended in cryopreservation medium before controlled-rate freezing and storage in liquid nitrogen until use. Cryopreserved PBMCs from HIV elite controllers were obtained from the Ragon Institute Clinical Data and Biobank (CIDB). All samples were collected and used in accordance with institutional guidelines.

### Tetramer staining

PBMCs were thawed at 37°C, diluted in pre-warmed R10 medium (RPMI 1640 (Sigma-Aldrich) supplemented with 10% FBS (Sigma-Aldrich), L-glutamine (Gibco), penicillin/streptomycin (Gibco), and HEPES (Gibco)), and treated with nuclease (Thermo Scientific) to reduce clumping. Cells were washed (300 × g, 10 min), resuspended, and pre-treated for 1 h at room temperature with either anti-CD94 blocking antibody (10 µg/mL; clone HP-3D9; BD Biosciences) plus dasatinib (50 nM; Selleck Chemicals), or dasatinib alone. Following incubation, cells were stained with HLA-E tetramers (final concentration of 20 nM) for 20 min at 4°C, followed by surface antibody staining for 30 min at 4°C. Cells were washed, fixed with Cytofix (BD Biosciences), and resuspended in FACS buffer prior to acquisition.

### Flow cytometric Immunophenotyping

Multiparametric immunophenotyping was performed on cryopreserved human PBMCs using a series of surface and intracellular staining panels designed to assess differentiation state, NK receptor expression, cytotoxic potential, and transcriptional profiles of tetramer-positive CD8⁺ T cell populations. All antibody panels were designed to minimize spectral overlap and empirically optimized prior to experimental use. Antibodies were titrated across a range of dilutions (1:25-1:200) in parallel staining conditions, and optimal concentrations were selected based on maximal resolution between positive and negative populations, assessed by separation of fluorescence intensity distributions rather than absolute signal intensity. Final panels were validated by re-staining with selected optimal dilutions to ensure compatibility across all markers within each panel.

### Surface staining

PBMCs were plated in 96-well plates and incubated with Fc receptor blocking reagent prior to staining. Where indicated, cells were pre-incubated with blocking antibodies targeting CD94/NKG2A/C (anti-CD94 blocking antibody; clone HP-3D9; BD Biosciences) to prevent tetramer-mediated non-TCR-directed binding. Cells were then stained with HLA-E*01:03 or HLA-A*02:01 tetramers (at a final concentration of 20 nM) at 4 °C, followed by incubation with surface antibody cocktails for 30 min at 4 °C protected from light. Following staining, cells were washed in FACS buffer and processed for either direct acquisition or downstream intracellular staining.

### Intracellular staining

For detection of intracellular molecules, cells were fixed and permeabilized using standard fixation/permeabilization buffers (BD Biosciences) following surface staining. Cells were incubated with antibodies against cytoplasmic molecules in permeabilization buffer (Thermofisher Scientific), followed by washing and fixation prior to acquisition. Intracellular staining of transcription factors was performed using a dedicated fixation and permeabilization system optimized for nuclear antigen detection. Following surface staining, cells were fixed and permeabilized using a transcription factor staining buffer kit (BD Biosciences), then incubated with antibodies targeting transcriptional regulators in permeabilization buffer. Cells were subsequently washed and resuspended in FACS buffer for acquisition.

### Activation assays

PBMCs were thawed and rested overnight at 37°C in complete R10 medium. Where indicated, cells were pre-treated with the IL-15-based superagonist, het-IL-15 (PBL Assay Science), for 48 h prior to assay. Cells were pre-treated with anti-CD94 blocking antibody (10 µg/mL) for 1 h, and anti-CD107a antibody (H4A3; BioLegend) was added immediately prior to stimulation. Cells were stimulated with fluorophore-conjugated HLA-E tetramers (10 nM final concentration) or PMA (positive control, eBioscience Cell Stimulation Cocktail) and incubated for 4 h at 37°C. Following this stimulatory incubation period, cells were stained with antibodies against CD3, CD8, CD69, and viability dye, fixed with Cytofix, and acquired on a flow cytometer.

### Flow cytometry

Data were acquired on a multicolor flow cytometer (BD FACS Symphony A5 analyzer) using single-stained compensation controls and fluorescence-minus-one (FMO) controls prior to analysis in FlowJo (v11.1.1). Lymphocytes were gated based on forward and side scatter, followed by singlets, live cells, CD3⁺CD8⁺ T cells, and tetramer-positive populations. Tetramer-positive populations were further analyzed for phenotypic and functional marker expression.

### Immunoregulatory *ex vivo* elimination assays

*Ex vivo* target cell elimination assays performed in this study were adapted from previously established methods^84^. Preparation of CD4^+^ T cell targets is as follows. Cryopreserved PBMCs were thawed and rested overnight at 37 °C in complete medium (RPMI 1640 supplemented with 10% fetal bovine serum, 1% L-glutamine, 1% penicillin–streptomycin, and HEPES). CD4⁺ T cells were isolated by negative selection using a magnetic bead–based enrichment kit (StemCell Technologies) according to the manufacturer’s instructions, in which unwanted cell populations are labeled with antibody complexes and removed using magnetic particles, yielding untouched CD4⁺ T cells. For non-specific activation, isolated CD4⁺ T cells were cultured on plates pre-coated with anti-CD3 antibody (2 µg⁻¹ in carbonate buffer; clone SK7; BioLegend) in non-tissue culture– treated plates to facilitate antibody immobilization. Cells were supplemented with complete medium containing IL-2 (10 ng ml⁻¹; PeproTech), and co-stimulated with soluble anti-CD28 antibody (4 µg ml⁻¹; clone CD28.2; BioLegend) for 72 h at 37 °C. Activation was confirmed by the presence of blast morphology via light microscopy prior to use. In parallel, non-activated CD4⁺ T cells were isolated from rested PBMCs on the day of assay setup using the same negative selection approach. Activated CD4⁺ T cells were labeled with CellTrace Violet (CTV; Invitrogen) according to the manufacturer’s instructions. Subsequently, both activated and non-activated CD4⁺ T cells were labeled with CellTrace Far Red (CTFR; Invitrogen) to distinguish total target populations. Briefly, cells were incubated with CTFR for 5 min at 37 °C, followed by quenching and washing in complete medium. Activated (CTV⁺CTFR⁺) and non-activated (CTV⁻CTFR⁺) CD4⁺ T cells were combined at a ratio of 30:70, respectively, to generate target populations for co-culture. HLA-E dependence was assessed by pre-treating activated CD4⁺ T cell targets with an anti-HLA-E blocking antibody (clone 3D12; 10–40 µg ml⁻¹; BioLegend), with blockade maintained throughout co-culture. Isotype-matched controls were included in parallel. Target CD4⁺ T cells were plated for co-culture at 3 × 10⁴–1 × 10⁵ cells per well (depending on availability) in TC-treated 96-well round-bottom plates.

Preparation of CD8^+^ T cell effectors is as follows. PBMCs were rested overnight prior to CD8⁺ T cell isolation. CD8⁺ T cells were isolated by negative selection using a magnetic bead–based enrichment kit (StemCell Technologies), yielding untouched effector cells for functional assays. Tetramer-based depletion controls were also performed via selective removal of CD94-blocked HLA-E/VL9-specific CD8⁺ T cells. For magnetic-based depletion, PBMCs were pre-blocked with anti-CD94 antibody to prevent non-TCR-mediated tetramer binding, followed by staining with APC-conjugated HLA-E/VL9, irrelevant HLA-A2/NV9 tetramers or no tetramer. Tetramer-positive cells were depleted using an anti-APC magnetic selection system (StemCell Technologies) according to the manufacturer’s protocol, including two sequential rounds of depletion to maximize depletion efficiency. For toxin-mediated depletion, HLA-E/VL9 tetramers were assembled using streptavidin conjugated to saporin (Advanced Targeting Systems), a ribosome-inactivating protein toxin that induces cell death upon internalization. In accordance with previously established methods^85^, PBMCs were incubated with saporin-conjugated tetramers (including HLA-E/VL9 tetramer, irrelevant HLA-A2*02:01/NV9 tetramer, or unconjugated saporin) for 2 h, followed by extensive washing and resting for up to 72 h prior to co-culture. Depletion efficiency was assessed by tetramer staining after a 6-day recovery period to allow for TCR re-expression. Mock depletion controls were performed in parallel using identical conditions without tetramer and with unconjugated saporin.

Freshly isolated, tetramer-depleted, or mock-depleted CD8⁺ T cells were added at defined effector-to-target (E:T) ratios of 0:1, 1:1, 2:1, 4:1, 8:1, and 16:1, with target-only wells included as controls. Co-cultures were established in a total volume of 200 µl per well in complete medium and incubated for 24 h at 37 °C. Following incubation, cells were harvested and stained with surface antibodies including viability dye to identify live target populations. Target cells were defined as live, singlet CD3⁺CD4⁺CTFR⁺ lymphocytes, and elimination was quantified as the reduction in the frequency of activated (CTV⁺CFTR^+^) relative to non-activated (CTV^-^ CTFR^+^) CD4^+^ T cells across increasing E:T ratios by flow cytometry. Specifically, percentage elimination was calculated as relative loss of activated targets in each E:T ratio compared with the target-only condition. For each E:T ratio, the frequency of activated targets among total targets was quantified and normalized to the frequency of activated targets in the target-only condition. Percentage elimination was calculated as:

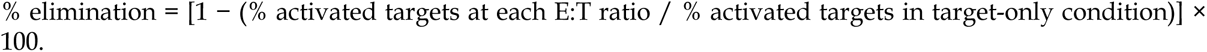

### Generation of VSV-G pseudotyped ΔEnv NL4-3 HIV virus

VSV-G-pseudotyped NL4-3 HIV particles were generated by transient transfection of HEK293T cells (ATCC). Cells were seeded in T225 flasks (Corning) and cultured to 60-80% confluency at the time of transfection. Plasmid DNA was diluted in Opti-MEM (Gibco), including 7.5 µg of VSV glycoprotein expression plasmid (pMD2.G; Addgene) and 64.5 µg of plasmid encoding ΔEnv-GFP NL4-3 HIV (NIH-AIDS Reagent Program, Cat# 11100 HIV-1 NL4-3 ΔEnv EGFP Reporter Vector). Polyethylenimine (PEI; Sigma-Aldrich) was added at a 3:1 PEI:DNA mass ratio, mixed gently, and incubated for 15 min at room temperature to allow formation of DNA-PEI complexes. Transfection mixtures were added dropwise to HEK293T cultures in complete growth medium without disturbing the cell monolayer. After 3 h at 37 °C, transfection medium was replaced with 30 ml pre-warmed complete DMEM(ATCC). Cells were then incubated for 72 h to allow viral production. Viral supernatants were harvested and clarified by centrifugation at 300 × g for 5 min to remove cellular debris, followed by filtration through a 0.45 µm (Millipore Sigma) syringe filter. Clarified viral supernatant was aliquoted and stored at −80 °C prior to use. Virus was used unconcentrated for downstream applications.

### *Ex vivo* HIV elimination assays

*Ex vivo* HIV elimination assays were performed with the same setup as the immunoregulatory elimination assays conducted in this study, except for the following modifications. CD4⁺ T cells were first isolated and activated using plate-bound anti-CD3 and soluble anti-CD28 antibodies for 72 h at 37 °C. Activated CD4⁺ T cells were then harvested and infected with GFP-encoding VSV-G-pseudotyped ΔEnv NL4-3 HIV pseudovirus by spinoculation. Briefly, cells were resuspended at 1 × 10⁶ cells ml⁻¹, virus was added at a 1:1 ratio (cells:virus; unconcentrated) with 8 ug/mL dextran, and plates were centrifuged for 2 h at 800 x g at 25 °C. Following spinoculation, cells were washed and resuspended in fresh culture medium supplemented with 10 ng/mL IL-2 and incubated for a further 72 h to allow productive infection. For elimination assays, infected CD4⁺ T cells were mixed with control CD4⁺ T cells that had been activated across identical timelines but not exposed to virus. In some experiments, infected cells were instead mixed with donor-matched, freshly isolated, non-infected, non-activated CD4⁺ T cells. All target cell populations were labeled with CTFR prior to co-culture. Productively infected cells were identified by CD4^low^ GFP^+^ expression by flow cytometry.

Target cells were co-cultured with autologous CD8⁺ T cells at increasing E:T ratios, as described above. Relative elimination was determined by comparing the frequency of CD4^low^ GFP⁺ (productively HIV-infected) cells to CD4^high^ GFP⁻ (non-infected HIV-negative) activated CD4⁺ T cell controls, as well as to HIV-negative non-activated CD4⁺ T cell targets. Specifically, for each E:T ratio, the frequency of HIV-infected target cells among total target cells was quantified and normalized to the frequency of HIV-infected target cells in the target-only condition. Percentage elimination was calculated as:

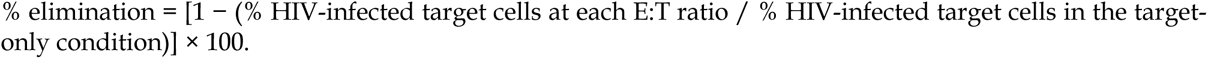

To assess HLA-E dependence, assays were performed in the presence of anti-HLA-E blocking antibody (clone 3D12; 10–40 µg ml⁻¹; BioLegend) or isotype control. In parallel, selective depletion of HLA-E/VL9 tetramer-positive CD8⁺ T cells was performed prior to co-culture, as described above.

### Single-cell TCR sequencing

Cryopreserved peripheral blood mononuclear cells (PBMCs) from eight donors were thawed and rested overnight at 37 °C with 5% CO₂ in R10 medium. Donors included two HIV-positive elite controllers, four HIV-negative CMV-seropositive persons with expanded E-sensor responses, and two CMV-seropositive individuals with expanded CMV-specific HLA-A02:01/NV9-specific responses. Elite controller samples were analyzed for HLA-E/VL9-specific CD8⁺ T cells, donor-matched immunodominant HIV peptide–HLA class I-specific responses, and unstained controls. HIV-negative donor samples were analyzed using one of three staining configurations: (i) HLA-E/VL9 and HLA-A02:01/NV9 tetramers with matched unstained controls (n = 1 donor), (ii) HLA-E/VL9 tetramers with unstained controls only (n = 3 donors), or (iii) HLA-A*02:01/NV9 tetramers with unstained controls only (n = 2 donors).

CD8⁺ T cells were enriched by magnetic negative selection using a custom antibody cocktail targeting CD14, CD19, and CD4 (StemCell Technologies). Enriched cells were incubated with anti-CD94 blocking antibody for 1 hour at room temperature. Cells were subsequently resuspended in monocyte blocker containing 1% dextran sulfate and pre-treated with 50 nM dasatinib (Selleck Chemicals) for 30 minutes at 37 °C to stabilize TCR– pHLA interactions. Cells were stained with peptide–HLA tetramers comprising HLA-E/VL9 or control peptide–HLA complexes conjugated to streptavidin-PE or streptavidin-APC following biotin quenching (77-fold molar excess). Following tetramer staining, cells were incubated with Fc receptor blocking reagent (TrueStain FcX; BioLegend) and stained with anti-CD3-BV510 (clone, BioLegend, dilution), anti-CD8-BV711 (clone, BioLegend, dilution), TotalSeq-C Human Universal Cocktail v2.0 (BioLegend), and sample-specific TotalSeq-C hashing antibodies (BioLegend; 1:200 dilution). Cells were washed using an HT2000 laminar flow cell washer (Curiox Biosystems) and resuspended in PBS containing 2% FBS and Sytox Green viability dye (Thermo Fisher Scientific).

Viable CD3⁺CD8⁺ tetramer-positive cells were sorted by fluorescence-activated cell sorting using a BD FACSAria Fusion. APC- and PE-tetramer double-positive populations were sorted and pooled within donors prior to loading onto Chromium GEM-X chips (10x Genomics) at approximately 30,000 cells per channel. Gene expression (GEX), antibody-derived tag (ADT), and T cell receptor (TCR) V(D)J libraries were generated using the Chromium GEM-X Single Cell 5′ platform according to the manufacturer’s instructions, including GEM generation, reverse transcription, cDNA amplification, and library construction. Libraries were sequenced on an Illumina NextSeq 2000 using a P3 100-cycle kit.

### Data analysis

Base calling and library level demultiplexing were performed using bcl2fastq2 v2.20.0.422. UMI quantitation was performed using the Cell Ranger (v10.0.0; 10x Genomics) multianalysis pipeline with the GRCh38 reference genome, and specifying “Gene Expression”, “Antibody Capture”, and “VDJ-T” feature types, and a custom feature reference including antibody derived tags and hashtags. Downstream analyses were performed in R (v4.3.1) using Seurat (v5.4.0.9008). Hashtag-based sample demultiplexing was performed using HTOdemux(), and only singlet cells (HTO_classification.global = “Singlet”) were retained for downstream analyses. Tetramer specificity assignments were used to define antigen-specific CD8⁺ T cell populations. TCR clonotypes were defined using productive TRB CDR3 amino acid sequences, and singleton clonotypes were excluded from clonality analyses, except for the case of calculation of clonal diversity. Clonal diversity was assessed using the inverse Simpson diversity index. Enriched clonotypes were identified by comparison with the corresponding bulk CD8⁺ T cell repertoire, and TCR gene usage and CDR3 sequence analyses were performed using productive paired TCR sequences.

All other data in this study were analyzed using FlowJo v11 (BD Biosciences), and statistical analyses and graph generation were performed using GraphPad Prism v11 (GraphPad Software). Diagrammatic figures were created using BioRender (BioRender.com).

## ACKNOWLEDGEMENTS

We thank all study participants for their invaluable contribution to this work. We additionally acknowledge the Ragon Institute flow cytometry core facilities staff for their support with experimental and technical aspects of the project. This work was supported by the Howard Hughes Medical Institute (HHMI). The mass spectrometry studies were additionally supported by NIH grant R01AI022553.

## AUTHOR CONTRIBUTIONS

L.C.W. conceptualized the study, designed and performed experiments, analyzed and interpreted data, prepared figures, and wrote the manuscript. B.D.W. supervised and funded the study, provided critical conceptual and technical development and interpretation throughout the project, and extensively edited and assisted in preparation of the manuscript. C.C. and X.G. contributed to protein expression, purification, refolding, monomer generation, tetramer staining, flow-based phenotyping, elimination assays, and single-cell T cell sequencing experiments. X.G. additionally performed activation assays. L.C.W. and O.L. performed mass spectrometry experiments; O.L. performed mass spectrometry data analysis and contributed to preparation and interpretation of the mass spectrometry data. B.D.B. and F.M.W. supervised the mass spectrometry studies and provided conceptual and technical guidance. D.R.C. provided detailed technical training and conceptual guidance throughout the study and assisted with TCR sequencing experiments. M.C. and M.E.B. provided conceptual and technical guidance throughout the project. H.W. assisted with single-cell T cell sequencing experiments, and J.M.U. performed downstream computational and bioinformatic analyses. E.C. provided technical guidance and training relating to cellular assays. A.O.C. provided guidance relating to T cell sequencing approaches and assisted with leukopak processing. P.A.B. assisted with recombinant protein production, protein refolding, and size-exclusion chromatography methodologies. All authors reviewed and approved the final manuscript.

## DATA AND CODE AVAILABILITY

Single-cell sequencing data generated in this study have been deposited in the NCBI Gene Expression Omnibus (GEO) prior to publication. The GRCh38 reference genome is available from NCBI GenBank (GCA_000001405.15).

