## Supplement for "Self-reactive human CD8^+^ T cells restricted by HLA-E mediate immunoregulatory and antiviral functions"

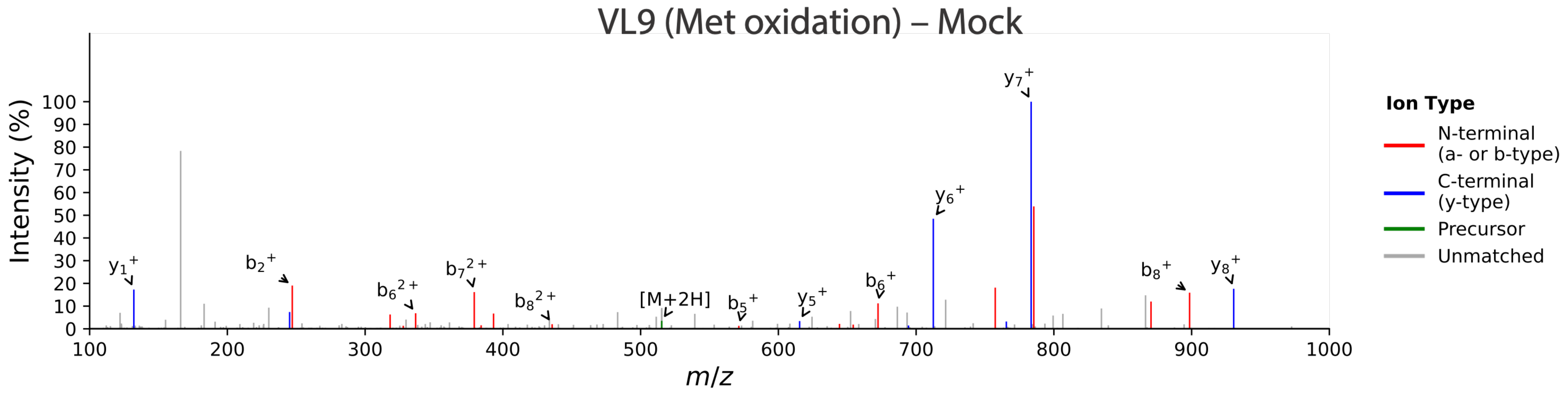


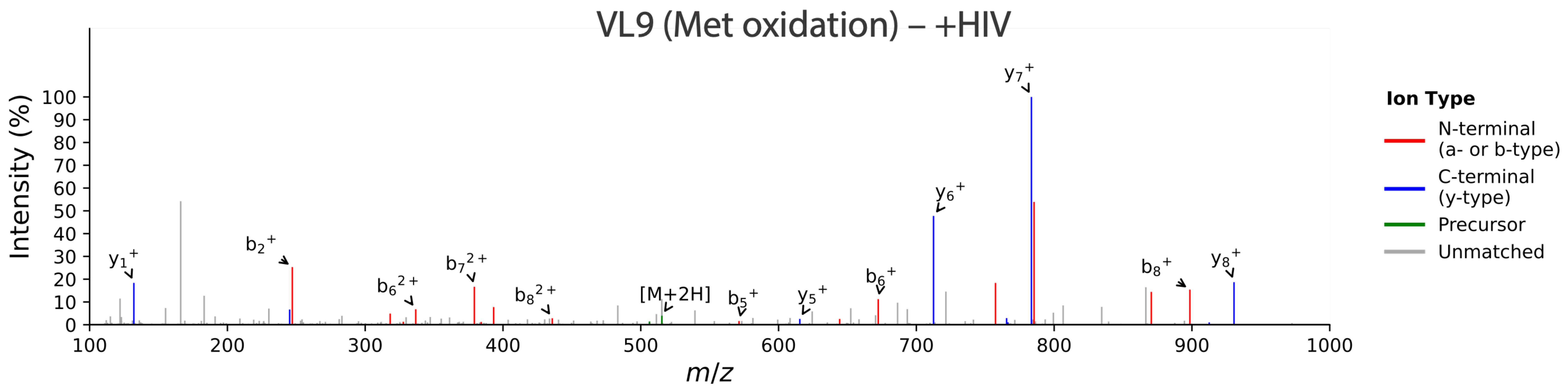


**Extended Data Figure 1. Representative MS/MS spectra demonstrating endogenous presentation of VL9 peptides following HIV infection.** Representative mass spectra of the endogenous VL9 peptide recovered following HLA-E immunoprecipitation from mock-infected (top) and HIV-infected (bottom) cells. Annotated b- and y-type fragment ions confirm peptide identity. Comparable fragmentation spectra demonstrate equally strong presentation of the endogenous oxidized VL9 peptide under both conditions.

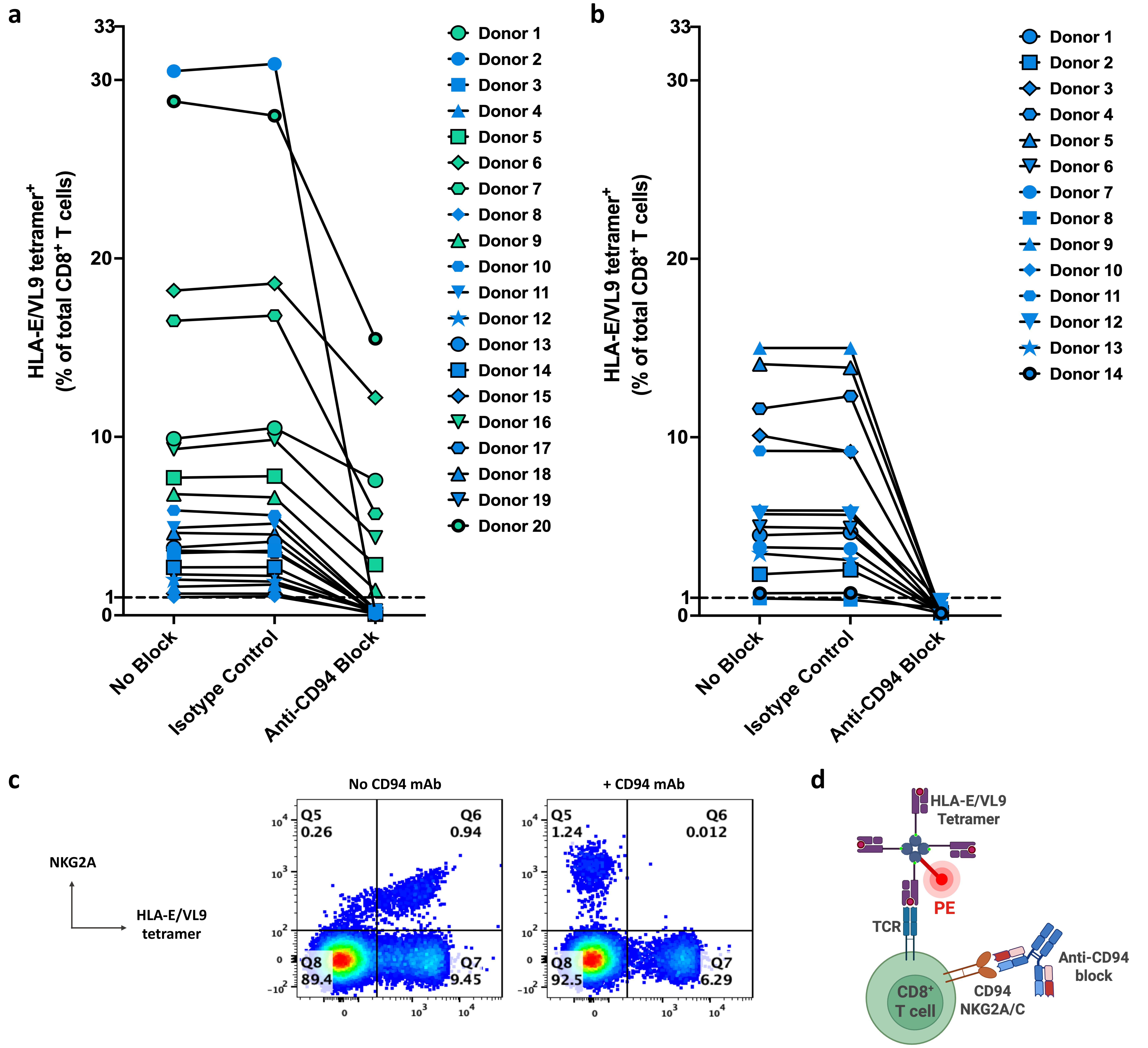


**Extended Data Figure 2. Validation of anti-CD94 antibody-mediated blockade.** (**a,b**) *Ex vivo* tetramer staining was performed on PBMC isolated from 20 healthy CMV seropositive (a) and 14 healthy CMV seronegative (b) donors. Cells were treated with an anti-CD94 blocking antibody, an isotype control antibody, or an absence of blocking antibody prior to staining with antibodies to CD3 and CD8α, in addition to HLA-E*01:03 tetramer refolded with the VMAPRTLIL peptide. Cells were gated on single, live, CD3/CD8α double-positive T lymphocytes, and HLA-E*01:03/VMAPRTLIL tetramer$^+$ cells are plotted as a percentage of total CD8$^+$ T cells. **(c)** PBMC isolated from CMV$^+$ Donor 16 were stained with antibodies to CD3, CD8α, and NKG2A, in addition to HLA-E*01:03 tetramer refolded with the VMAPRTLIL peptide. Prior to staining, cells were incubated with and without an anti-CD94 blocking antibody. Quadrants show the proportion of NKG2A$^+$ and HLA-E/VMAPRTLIL tetramer$^+$ cells after gating on single, live, CD3/CD8α double-positive T lymphocytes. **(d)** Schematic illustrating the CD94 blockade strategy. Anti-CD94 antibody prevents HLA-E/VL9 tetramer binding to CD94/NKG2A and CD94/NKG2C receptors, enabling selective detection of HLA-E/VL9-specific TCR-mediated tetramer staining.

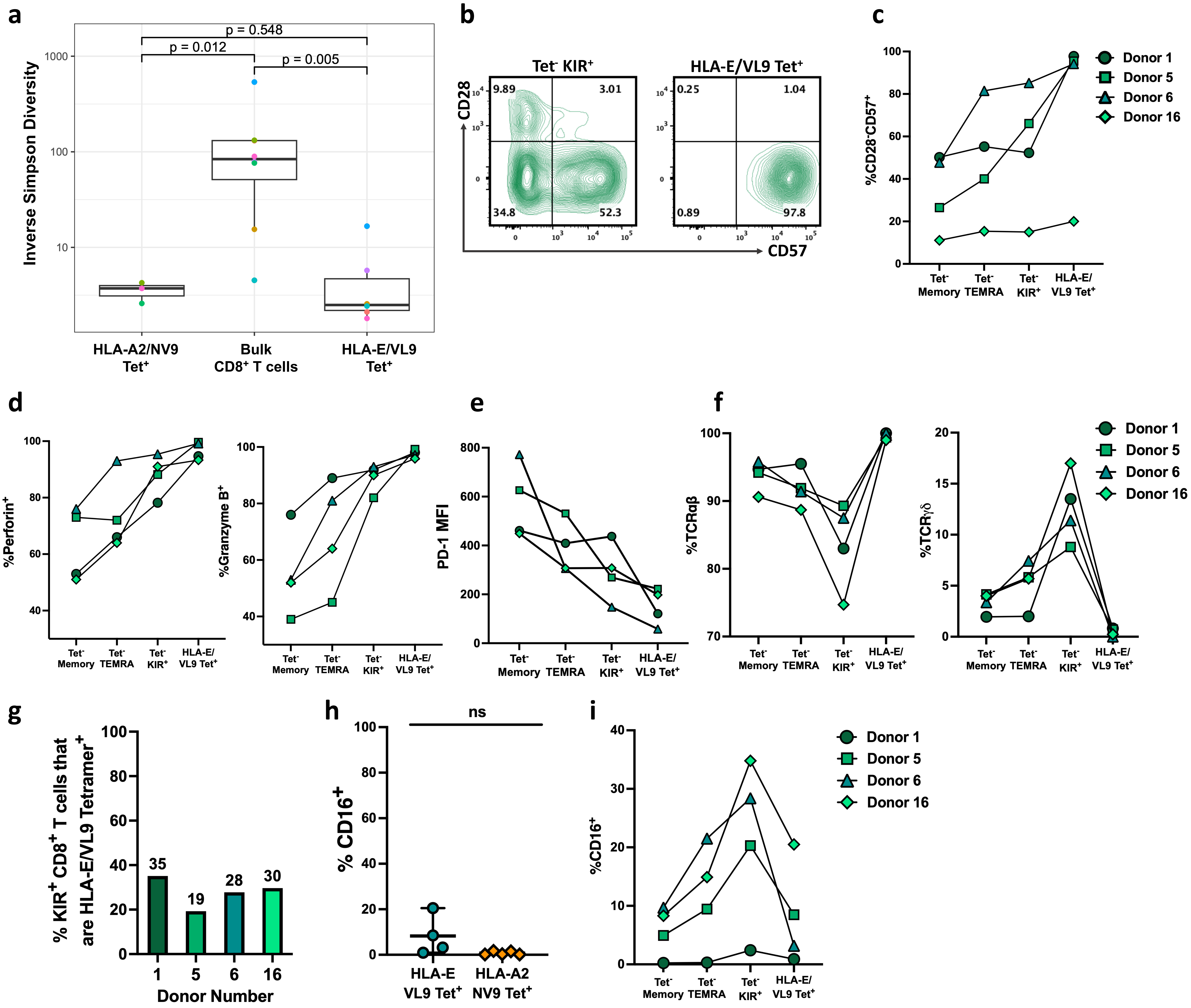


**Extended Data Figure 3. Phenotypic and clonotypic characterization of E-sensors. (a)** Clonal TCR diversity was quantified using the inverse Simpson diversity index for HLA-E/VL9 tetramer-positive CD8⁺ T cells (n = 6 donors), HLA-A*02:01/NV9 tetramer-positive *CD8⁺ T* cells *(n = 3 CMV-seropositive HLA-A**02:01⁺ donors), and donor-matched bulk CD8⁺ T cells. Each point represents an individual donor. Statistical significance was assessed using two-sided Wilcoxon rank-sum tests. **(b)** Representative flow cytometry plots using a CMV-seropositive donor with an expanded (>1% of CD8⁺ T cells) E-sensor population. CD28 and CD57 expression are shown for HLA-E/VL9 tetramer-negative KIR⁺ CD8⁺ T cells and HLA-E/VL9 tetramer-positive CD8⁺ T cells. Frequencies of cells within each quadrant are indicated. **(c)** Summary of the frequency of CD28⁻CD57⁺ cells among tetramer-negative CD8⁺ T cell memory, tetramer-negative TEMRA, tetramer-negative KIR⁺ (KIR2D⁺ and/or KIR3D⁺), and HLA-E/VL9 tetramer-positive CD8⁺ T cells from four healthy CMV-seropositive donors (D1, D5, D6 and D16). **(d)** Summary of perforin (left) and granzyme B (right) expression among the CD8⁺ T cell populations shown in b. **(e)** Summary of PD-1 expression among the CD8⁺ T cell populations shown in b. **(f)** Summary of αβ TCR (left) and γδ TCR (right) expression among the CD8⁺ T cell populations shown in b. **(g)** Frequency of HLA-E/VL9 tetramer staining among total KIR-expressing (KIR2D and/or KIR3D) CD8⁺ T cells. **(h)** Frequency of CD16⁺ cells among HLA-A*02:01/NV9 (an immunodominant CMV peptide) tetramer-positive and HLA-E/VL9 tetramer-positive CD8⁺ T cells from CMV seropositive donors. Lines indicate medians; the line connecting groups indicates a lack of statistically significant differences by two-tailed t-tests. **(i)** Summary of CD16 expression among the CD8⁺ T cell populations shown in b.

| Donor | CMV Status | Age | Sex | Ethnicity | MHC Class Ia and VL9 status | | | | | |
|---|---|---|---|---|---|---|---|---|---|---|
| 1 | *POS* | 42 | F | CAU | A*24:02<br>VMAPRTLVL | A*26:01<br>VMAPRTLVL | B*18:01<br>VTAPRTLLL | B*49:01<br>VTAPRTVLL | C*07:01<br>VMAPRALLL | C*07:01<br>VMAPRALLL |
| 2 | *POS* | 60 | F | CAU | A*01:01<br>VMAPRTLLL | A*24:02<br>VMAPRTLVL | B*07:02<br>VMAPRTVLL | B*08:01<br>VMAPRTVLL | C*07:01<br>VMAPRALLL | C*07:02<br>VMAPRALLL |
| 3 | *POS* | 22 | F | MIX | A*03:01<br>VMAPRTLLL | A*23:01<br>VMAPRTLVL | B*08:01<br>VMAPRTVLL | B*58:01<br>VTAPRTVLL | C*07:01<br>VMAPRALLL | C*07:01<br>VMAPRALLL |
| 4 | *POS* | 47 | M | UNK | A*02:01<br>VMAPRTLVL | A*02:01<br>VMAPRTLVL | B*40:02<br>VTAPRTLLL | B*55:02<br>VTAPRTLLL | C*02:02<br>VMAPRTLLL | C*15:02<br>VMAPRTLLL |
| 5 | *POS* | 24 | M | ASN | A*24:02<br>VMAPRTLVL | A*24:02<br>VMAPRTLVL | B*07:02<br>VMAPRTVLL | B*44:03<br>VTAPRTLLL | C*07:01<br>VMAPRALLL | C*07:02<br>VMAPRALLL |
| 6 | *POS* | 41 | M | CAU | A*02:01<br>VMAPRTLVL | A*26:01<br>VMAPRTLVL | B*27:05<br>VTAPRTLLL | B*49:01<br>VTAPRTVLL | C*02:02<br>VMAPRTLLL | C*07:01<br>VMAPRALLL |
| 7 | *POS* | 39 | M | CAU | A*02:01<br>VMAPRTLVL | A*24:02<br>VMAPRTLVL | B*18:01<br>VTAPRTLLL | B*27:05<br>VTAPRTLLL | C*02:02<br>VMAPRTLLL | C*07:01<br>VMAPRALLL |
| 8 | *POS* | 36 | M | AFA | A*68:02<br>VMAPRTLVL | A*68:02<br>VMAPRTLVL | B*15:03<br>VTAPRTVLL | B*42:02<br>VMAPRTVLL | C*02:10<br>VMAPRTLLL | C*17:01<br>VMAPQALLL |
| 9 | *POS* | 37 | F | CAU | A*02:01<br>VMAPRTLVL | A*26:01<br>VMAPRTLVL | B*27:05<br>VTAPRTLLL | B*49:01<br>VTAPRTVLL | C*02:02<br>VMAPRTLLL | C*07:01<br>VMAPRALLL |
| 10 | *POS* | 42 | M | HSP | A*29:02<br>VMAPRTLLL | A*68:01<br>VMAPRTLVL | B*44:03<br>VTAPRTLLL | B*52:01<br>VTAPRTVLL | C*03:04<br>VMAPRTLIL | C*16:01<br>VMAPRTLIL |
| 11 | *POS* | 48 | M | CAU | A*02:01<br>VMAPRTLVL | A*03:01<br>VMAPRTLLL | B*07:02<br>VMAPRTVLL | B*37:01<br>VTAPRTLLL | C*06:02<br>VMAPRTLIL | C*07:02<br>VMAPRALLL |
| 12 | *POS* | 46 | M | CAU | A*02:01<br>VMAPRTLVL | A*03:01<br>VMAPRTLLL | B*08:01<br>VMAPRTVLL | B*15:01<br>VTAPRTVLL | C*03:04<br>VMAPRTLIL | C*07:01<br>VMAPRALLL |
| 13 | *POS* | 52 | M | CAU | A*24:02<br>VMAPRTLVL | A*32:01<br>VMAPRTLLL | B*44:02<br>VTAPRTLLL | B*51:09<br>VTAPRTVLL | C*01:02<br>VMAPRTLIL | C*07:04<br>VMAPRALLL |
| 14 | *POS* | 57 | F | CAU | A*03:01<br>VMAPRTLLL | A*24:02<br>VMAPRTLVL | B*15:01<br>VTAPRTVLL | B*35:08<br>VTAPRTVLL | C*03:03<br>VMAPRTLIL | C*04:01<br>VMAPRTLIL |
| 15 | *POS* | 46 | F | AFA | A*03:01<br>VMAPRTLLL | A*24:02<br>VMAPRTLVL | B*15:10<br>VTAPRTVLL | B*44:02<br>VTAPRTLLL | C*04:01<br>VMAPRTLIL | C*07:02<br>VMAPRALLL |
| 16 | *POS* | 40 | F | CAU | A*01:01<br>VMAPRTLLL | A*02:01<br>VMAPRTLVL | B*40:01<br>VTAPRTVLL | B*44:02<br>VTAPRTLLL | C*03:04<br>VMAPRTLIL | C*07:04<br>VMAPRALLL |
| 17 | *POS* | 37 | M | WHT | A*02:01<br>VMAPRTLVL | A*11:01<br>VMAPRTLLL | B*39:24<br>VMAPRTVLL | B*52:01<br>VTAPRTVLL | C*07:01<br>VMAPRALLL | C*12:02<br>VMAPRTLIL |
| 18 | *POS* | 30 | M | HSP | A*02:01<br>VMAPRTLVL | A*24:02<br>VMAPRTLVL | B*39:06<br>VMAPRTVLL | B*40:02<br>VTAPRTLLL | C*03:04<br>VMAPRTLIL | C*07:02<br>VMAPRALLL |
| 19 | *POS* | 21 | M | ASN | A*02:11<br>VMAPRTLVL | A*03:01<br>VMAPRTLLL | B*52:01<br>VTAPRTVLL | B*55:01<br>VTAPRTLLL | C*01:02<br>VMAPRTLIL | C*12:02<br>VMAPRTLIL |
| 20 | *POS* | 31 | F | CAU | A*02:01<br>VMAPRTLVL | A*31:01<br>VMAPRTLLL | B*15:01<br>VTAPRTVLL | B*44:02<br>VTAPRTLLL | C*03:03<br>VMAPRTLIL | C*05:01<br>VMAPRTLIL |

**Extended Data Table 1. Characteristics of healthy CMV-seropositive donors.** Donor demographics, classical HLA class I genotypes, and corresponding HLA class I signal peptide-derived VL9 sequences for healthy CMV-seropositive, HIV-negative donors screened for E-sensor responses in this study. Donors were selected based on previously reported genetic associations with HLA-E-restricted VL9-specific $CD8^+$ T cell expansions.

| Donor | CMV Status | Age | Sex | Ethnicity | MHC Class Ia and VL9 status | | | | | |
|---|---|---|---|---|---|---|---|---|---|---|
| 1 | *NEG* | 62 | M | CAU | **A*03:01**<br>VMAPRTLLL | **A*24:02**<br>VMAPRTLVL | **B*07:02**<br>VMAPRTVLL | **B*27:05**<br>VTAPRTLLL | **C*02:02**<br>VMAPRTLLL | **C*07:02**<br>VMAPRALLL |
| 2 | *NEG* | 45 | F | CAU | **A*01:01**<br>VMAPRTLLL | **A*02:01**<br>VMAPRTLVL | **B*07:02**<br>VMAPRTVLL | **B*08:01**<br>VMAPRTVLL | **C*07:01**<br>VMAPRALLL | **C*07:02**<br>VMAPRALLL |
| 3 | *NEG* | 58 | M | ASN | **A*02:11**<br>VMAPRTLVL | **A*31:01**<br>VMAPRTLLL | **B*07:02**<br>VMAPRTVLL | **B*40:06**<br>VTAPRTLLL | **C*07:02**<br>VMAPRALLL | **C*15:02**<br>VMAPRTLLL |
| 4 | *NEG* | 54 | M | CAU | **A*24:02**<br>VMAPRTLVL | **A*32:01**<br>VMAPRTLLL | **B*55:01**<br>VTAPRTLLL | **B*56:01**<br>VTAPRTLLL | **C*01:02**<br>VMAPRTLIL | **C*03:03**<br>VMAPRTLIL |
| 5 | *NEG* | 45 | M | CAU | **A*01:01**<br>VMAPRTLLL | **A*02:01**<br>VMAPRTLVL | **B*07:02**<br>VMAPRTVLL | **B*35:03**<br>VTAPRTVLL | **C*04:01**<br>VMAPRTLIL | **C*07:02**<br>VMAPRALLL |
| 6 | *NEG* | 55 | F | CAU | **A*02:01**<br>VMAPRTLVL | **A*03:01**<br>VMAPRTLLL | **B*13:02**<br>VTAPRTLLL | **B*44:02**<br>VTAPRTLLL | **C*05:01**<br>VMAPRTLIL | **C*06:02**<br>VMAPRTLIL |
| 7 | *NEG* | 53 | M | CAU | **A*02:01**<br>VMAPRTLVL | **A*68:01**<br>VMAPRTLVL | **B*35:03**<br>VTAPRTVLL | **B*44:02**<br>VTAPRTLLL | **C*04:01**<br>VMAPRTLIL | **C*05:01**<br>VMAPRTLIL |
| 8 | *NEG* | 29 | M | MIX | **A*30:01**<br>VMAPRTLLL | **A*68:01**<br>VMAPRTLVL | **B*07:02**<br>VMAPRTVLL | **B*58:02**<br>VTAPRTVLL | **C*06:02**<br>VMAPRTLIL | **C*07:02**<br>VMAPRALLL |
| 9 | *NEG* | 45 | M | CAU | **A*01:01**<br>VMAPRTLLL | **A*02:01**<br>VMAPRTLVL | **B*07:02**<br>VMAPRTVLL | **B*35:03**<br>VTAPRTVLL | **C*04:01**<br>VMAPRTLIL | **C*07:02**<br>VMAPRALLL |
| 10 | *NEG* | 18 | F | CAU | **A*02:01**<br>VMAPRTLVL | **A*30:01**<br>VMAPRTLLL | **B*13:02**<br>VTAPRTLLL | **B*27:05**<br>VTAPRTLLL | **C*02:02**<br>VMAPRTLLL | **C*06:02**<br>VMAPRTLIL |
| 11 | *NEG* | 52 | M | CAU | **A*02:01**<br>VMAPRTLVL | **A*68:01**<br>VMAPRTLVL | **B*35:03**<br>VTAPRTVLL | **B*44:02**<br>VTAPRTLLL | **C*04:01**<br>VMAPRTLIL | **C*05:01**<br>VMAPRTLIL |
| 12 | *NEG* | 29 | F | CAU | **A*02:01**<br>VMAPRTLVL | **A*66:01**<br>VMAPRTLVL | **B*41:02**<br>VTAPRTVLL | **B*44:02**<br>VTAPRTLLL | **C*05:01**<br>VMAPRTLIL | **C*17:01**<br>VMAPQALLL |
| 13 | *NEG* | 50 | M | CAU | **A*02:01**<br>VMAPRTLVL | **A*68:02**<br>VMAPRTLVL | **B*14:02**<br>VMAPRTVLL | **B*15:01**<br>VTAPRTVLL | **C*03:04**<br>VMAPRTLIL | **C*08:02**<br>VMAPRTLIL |
| 14 | *NEG* | 20 | M | HSP | **A*02:01**<br>VMAPRTLVL | **A*23:05**<br>VMAPRTLVL | **B*07:02**<br>VMAPRTVLL | **B*44:02**<br>VTAPRTLLL | **C*06:02**<br>VMAPRTLIL | **C*07:02**<br>VMAPRALLL |

**Extended Data Table 2. Characteristics of healthy CMV-seronegative donors.** Donor demographics, classical HLA class I genotypes, and corresponding HLA class I signal peptide-derived VL9 sequences for healthy CMV-seronegative, HIV-negative donors screened for expanded E-sensor populations in this study. Donors were selected based on previously reported genetic associations with HLA-E-restricted VL9-specific $CD8^{+}$ T cell expansions.
